# Direct and indirect regulation of β-glucocerebrosidase by the transcription factors *USF2* and *ONECUT2*

**DOI:** 10.1101/2024.04.28.591482

**Authors:** Kathi Ging, Lukas Frick, Johannes Schlachetzki, Andrea Armani, Yanping Zhu, Pierre-André Gilormini, Ana Marques, Ashutosh Dhingra, Desirée Böck, Matthew Deen, Xi Chen, Tetiana Serdiuk, Chiara Trevisan, Stefano Sellitto, Claudio Pisano, Christopher K Glass, Peter Heutink, Jiang-An Yin, David J Vocadlo, Adriano Aguzzi

## Abstract

Mutations in the *GBA* gene, which encodes the lysosomal enzyme β-glucocerebrosidase (GCase), are the most prevalent genetic susceptibility factor for Parkinson’s disease (PD). However, only approximately 20% of carriers develop the disease, suggesting the presence of genetic modifiers influencing the risk of developing PD in the presence of *GBA* mutations. Here we screened 1,634 human transcription factors (TFs) for their effect on GCase activity in cell lysates of the human glioblastoma line LN-229, into which we introduced the pathogenic *GBA* L444P variant via adenine base editing. Using a novel arrayed CRISPR activation library, we uncovered 11 TFs as regulators of GCase activity. Among these, activation of *MITF* and *TFEC* increased lysosomal GCase activity in live cells, while activation of *ONECUT2* and *USF2* decreased it. Conversely, ablating USF2 increased *GBA* mRNA and led to enhanced levels of GCase protein and activity. While MITF, TFEC, and USF2 affected *GBA* transcription, ONECUT2 was found to control GCase trafficking by modulating the guanine exchange factors PLEKHG4 and PLEKHG4B. Hence, our study provides a systematic approach to identifying modulators of GCase activity, expands the transcriptional landscape of *GBA* regulation, and deepens our understanding of the mechanisms involved in influencing GCase activity.

## Introduction

Parkinson’s disease (PD) is a progressive neurodegenerative disorder characterized by motor symptoms comprising bradykinesia, rigidity, postural instability, and tremor, as well as non-motor features, including depression and cognitive impairment (1). *GBA* mutations have emerged as a major genetic susceptibility factor for PD development (2). *GBA* encodes the lysosomal enzyme β- glucocerebrosidase (GCase), which catalyzes the hydrolysis of glucosylceramide (GlcCer) and glucosylsphingosine (GlcSph) into ceramide, sphingosine, and glucose. Thus, GCase is crucially involved in the metabolism of glycosphingolipids (GSL), which play central roles in growth regulation, cell migration, apoptosis, and inflammatory responses, among other processes (3).

The association between *GBA* mutations and PD was first identified in individuals with Gaucher disease (GD), the most common lysosomal storage disorder, caused by homozygous mutations in *GBA*. It then emerged that not only individuals with GD but also their relatives carrying heterozygous *GBA* mutations have an elevated risk of developing PD. This observation established *GBA* variants as the most common genetic susceptibility factor for PD, with a prevalence of 5-10% in PD patients (2, 4–7). Individuals with PD associated with *GBA* mutations exhibit an earlier age of disease onset and show greater cognitive impairment than non-mutation carriers (2, 5). Further strengthening the link between GCase and PD, the enzymatic activity of GCase is reduced in PD patients even in the absence of mutations within *GBA* (8). Thus, *GBA* appears to play a key role in the development and progression of PD.

Only about 20% of individuals with disease-associated *GBA* mutations will develop PD. This modest penetrance, along with the clustering of PD in families carrying *GBA* variants and phenotypic heterogeneity in GD symptoms, suggests the involvement of genetic modifiers influencing GCase activity, disease risk, and severity (4). Genome-wide association studies (GWAS) have identified *BIN1*, *SNCA*, *TMEM175*, and *CTSB* as candidate loci influencing PD risk and progression in the context of *GBA* mutations. However, their impact on GCase function is unknown (9). In addition, these candidate loci likely represent only a fraction of the genetic modifiers associated with disease risk, as low-prevalence single nucleotide variants (SNVs) may go undetected in GWAS (10).

Since transcription factors (TFs) often serve as "master regulators" orchestrating entire pathways and the expression of numerous genes, they represent attractive targets for identifying comprehensive networks involved in the regulation of processes of interest (11). Like many other lysosomal genes, the *GBA* promoter harbors two Coordinated Lysosomal Expression and Regulation (CLEAR) motifs, DNA sequences that are recognized by Transcription Factor EB (TFEB). This nuclear-lysosomal axis enables cells to fine-tune their metabolism in response to environmental changes, such as nutrient availability (12, 13). Interestingly, the presence of evolutionarily conserved sequences within the 5′ and 3′ non- coding regions of *GBA* suggest the existence of additional TF binding sites affording control of GCase expression by other, unidentified TFs (14). Yet, no TF has emerged from GWAS studies focused on PD susceptibility.

CRISPR-based screens offer a powerful means of systematically identifying genes regulating biological processes such as GCase activity. Unlike GWAS, which are correlative, CRISPR screens enable the discovery of causal relationships (15–17). Toward this goal, we recently generated an arrayed CRISPR activation (CRISPRa) library termed T.gonfio that targets each protein-coding gene with four non-overlapping guide RNAs (18). Using glioblastoma cells base-edited to harbor a pathogenic *GBA* mutation, we individually activated 1,634 human TFs. By assaying GCase in cell lysates, we identified 29 TFs modulating GCase activity including MITF and TFEC, two members of the MiT/TFE family of TFs to which TFEB and TFE3 also belong. *MITF* and *TFEC* increased whereas *USF2* and *ONECUT2* decreased lysosomal GCase activity, respectively. ONECUT2 activation did not regulate *GBA* transcript levels but influenced the intracellular trafficking of GCase. These findings expand the transcriptional landscape of *GBA* regulation, deepening our understanding of the mechanisms involved in modulating GCase activity in the context of *GBA* mutations.

## Results

### A cellular model system to study L444P mutant GCase activity

Large-scale genetic screens require scalable, easy-to-manipulate model systems. We selected the human glioblastoma line LN-229 which, in comparison with other cancer cell lines, possesses intermediate endogenous GCase activity and expression levels (19). As the detection of enhancers of mutant GCase might be clinically most relevant, we introduced the pathogenic L444P variant (NM_000157.4:c.1448T>C) into *GBA* (Fig. 1A) using the adenine base editor ABE8e (20). The L444P mutation results in misfolding of GCase that leads to recognition by the cellular quality-control system and proteasomal degradation. This mutation is strongly associated with the development of PD and causes a severe neurological syndrome in homozygous individuals (21–23).

**Figure 1.**
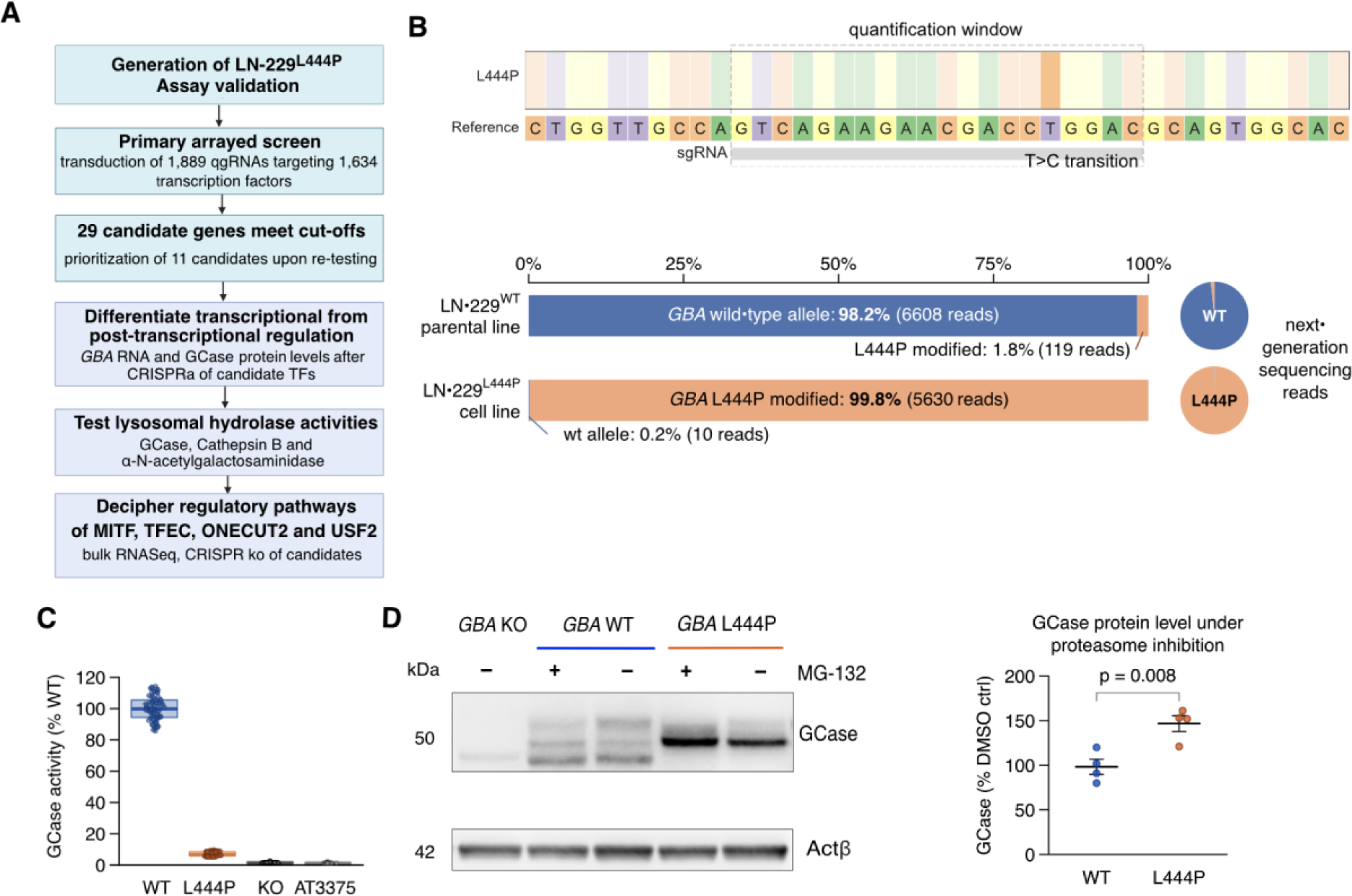
A forward-genetics screen for GCase activity modulators. A. Study workflow. B. Read frequencies (next-generation sequencing) of the desired T>C point mutation in the base-edited LN-229^L444P^ (orange) and its parental LN-229^WT^ (blue) line. C. GCase enzymatic activity in cell lysates of LN-229^WT^ (WT), LN-229^L444P^ (L444P), and GBA-ablated (KO) LN-229 cells. 30 wells/genotype, two independent experiments. AT3375: GCase-selective inhibitor. D. Immunoblot (left) and quantification (right) of cellular GCase protein in LN-229ΔGBA (KO), LN-229^L444P^ (L444P), and LN-229^WT^ (WT) after treatment with the proteasome inhibitor MG-132 (+) or with DMSO vehicle (−). N = 4 experiments, unpaired two-samples t-test.

Next-generation sequencing using primers that selectively amplify *GBA* but not its highly homologous pseudogene *GBAP1* (24) confirmed the presence of the L444P mutation in all alleles of the polyploid LN-229 line (Fig. 1B). A single-cell derived edited clone (henceforth referred to as LN-229^L444P^) was used for all subsequent experiments. GCase activity in LN-229^L444P^ lysates was reduced to 7% of the parental line expressing wild-type GCase (LN-229^WT^) (Fig. 1C) when measured with a lysate-based assay employing the artificial substrate 4-methylumbelliferyl-β-D-glucopyranoside (4MU-Glc). This is similar to the residual enzymatic activity reported in fibroblasts from patients with a homozygous L444P mutation (23). Treatment of LN-229^L444P^ cells with the proteasome inhibitor MG-132 increased GCase protein levels to 145% and GCase activity in cell lysates to 139% of DMSO-treated control cells, whereas GCase protein levels and activity remained unchanged in LN-229^WT^ cells (Fig. 1D and Supplementary Fig. 1A). These results support successful engineering of the *GBA* gene to introduce the clinically relevant L444P mutation within this clonal line.

One important caveat of lysate-based assays employing artificial substrates is hydrolysis by human β- glucosidases other than GCase, such as GBA2 and GBA3 (25–28). To ascertain the specificity of our assay for GCase activity in LN-229 cells, we conducted dose-response experiments in cell lysates using three inhibitors: conduritol-𝛽-epoxide (CBE), which inhibits glycosidases beyond GCase at higher concentrations (29), the highly selective GCase inhibitor AT3375 (27), and the GBA2-selective inhibitor miglustat (30).

Treatment of LN-229^WT^ cells with CBE and AT3375 at commonly employed working concentrations (300 µM and 10 µM, respectively) reduced the fluorescent signal to <5% of that seen in DMSO-treated cells, whereas miglustat caused no decrease (Supplementary Fig. 1B-D). To further confirm specificity of the assay, we generated LN-229 cells in which *GBA* was ablated (LN-229^ΔGBA^) by applying sgRNAs targeting the catalytic domain of GCase. In LN-229^ΔGBA^ lysates, the turnover of 4MU-Glc was reduced to 1.5% of the levels observed in LN-229^WT^ cells (Fig. 1C). Collectively, these data confirmed premature proteasomal degradation of the L444P mutant GCase in engineered LN-229^L444P^ cells. Additionally, dose-response curves showed that 4MU-Glc was predominantly hydrolyzed by GCase in LN-229 cells, eliminating the need for a subtractive assay using chemical inhibitors to correct for the contribution of other β-glucosidases (25). This led us to conclude that the LN-229^L444P^ cell line is suitable for identifying modulators of L444P mutant GCase activity using a lysate-based assay.

### An arrayed CRISPRa screen identifies 11 TFs as modulators of mutant GCase activity

*To identify TFs regulating GCase activity, we conducted an arrayed CRISPRa screen using the T.gonfio library* (*18*)*. To achieve high on-target activity, each protein-coding gene is targeted by four non-overlapping sgRNAs referred to as quadruple-guide RNAs (qgRNAs). The qgRNA plasmids were designed to tolerate human DNA polymorphisms and so maximize their versatility. In addition, each major transcription start site (TSS) was targeted individually to account for potential divergence in TSS activities among different cell types. We then performed a high-throughput arrayed transcriptional activation (CRISPRa) screen (Fig. 2A). LN-229^L444P^ cells stably expressing dCas9-VPR were seeded and transduced with lentiviruses carrying 1,889 qgRNAs targeting 1,634 human TFs* (*11*) *in three replicate 384-well plates, resulting in 6,729 transductions (including controls) with 171 TFs having more than one TSS. Five days post-transduction, GCase activity was measured on two replicate plates in cell lysates using 4MU-Glc as the substrate. On the third replicate plate, cell viability was assessed using the CellTiter-Glo assay (Supplementary* Fig. 2A).

**Fig. 2.**
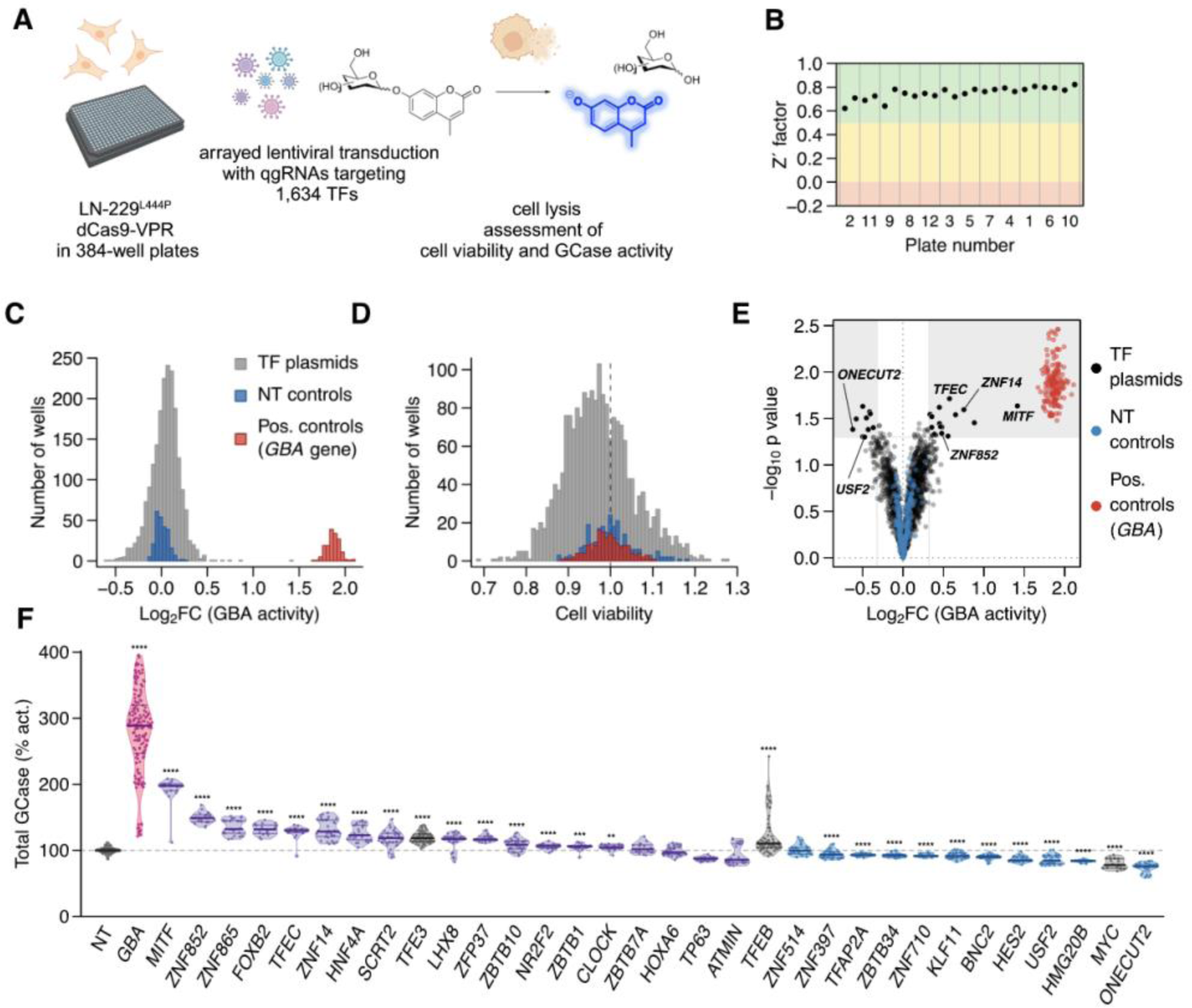
Arrayed CRISPR activation screen of human TFs in LN-229^L444P^ cells. A. Overview of the screening workflow. B. Z’ factor of each assay plate reporting on the separation between cells exposed to qgRNA lentiviruses targeting *GBA* (positive controls) vs. nontargeting (NT) qgRNA (negative controls). Plates are ordered in ascending order of Z’ factor of the replicate plates, with plate numbers (order of pipetting) provided on the x-axis. C-D. Histograms of enzymatic activity (C) and viability (D) of LN-229^L444P^ cells treated with TF-activating (grey), *GBA*-specific (red), and nontargeting qgRNAs (blue). GCase activity was normalized to wells infected with NT controls. E. Changes in GCase activity (abscissa) and ps (ordinate) after activation of designated transcription factors. F. Re-testing of 29 candidates with a higher number of replicates. Grey: NT controls and TFs previously found to regulate GCase expression (*TFE3*, *TFEB*, and *MYC*). Purple and blue: TFs increasing and decreasing GCase activity in the primary screen, respectively. Red: positive controls (*GBA*-activated cells). N ≥10 transductions in ≥2 independent experiments; unpaired t-test. Asterisks: ps (here and henceforth). **** p<0.0001; *** p<0.001; ** p<0.01; *p<0.05

Positive (*GBA*-activated) and non-targeting control (NT ctrl) qgRNAs yielded distinct signals with no overlap in fluorescence intensities, resulting in a Z′-factor >0.5 on all plates (31) (Fig. 2B-C). The coefficient of variation between duplicate samples (R^2^) was 0.71, demonstrating a high degree of technical reproducibility (Supplementary Fig. 2D). No major changes in cell viability were observed after lentiviral transduction and activation of TFs (Fig. 2D). Except for a minor plate gradient towards lower values around well A24 (Supplementary Fig. 2B), the quality measurements confirmed the robustness of the primary screen, instilling confidence in our selection of candidate TFs for further investigation.

Clinical observations in GD patients undergoing enzyme replacement therapy suggest that even small increases in GCase activity (≈10%) can be clinically meaningful (32). Hence, using the log_2_-transformed fold change (log_2_FC) fluorescence intensity values, we applied the following cut-off criteria for candidate selection: (1) a p <0.05 computed from two experimental replicates and (2) a mean log_2_FC > 0.32 or < −0.32 for up- and downregulators of GCase activity, respectively. A log_2_FC of ±0.32 resulted in 4.5 standard deviations (SD) from the mean fluorescence values of the NT controls. None of the 249 NT controls met these cut-off criteria. These selection criteria resulted in the nomination of 29 TFs (18 up- and 11 down-regulators of enzymatic activity) as candidate regulators of GCase activity (Fig. 2E). To prioritize the strongest candidates and minimize false-positive hits, we retested qgRNAs targeting those 29 candidate TFs in ≥2 independent experiments; each TF was tested with ≥10 replicates. 11 of the 29 candidates (38%), namely *MITF*, *ZNF852*, *ZNF865*, *FOXB2*, *TFEC*, *ZNF14*, *HNF4A*, *SCRT2*, *USF2*, *HMG20B*, *ONECUT2* exhibited changes in GCase activity equal to or greater than those observed following CRISPRa of *TFE3* (median log_2_FC = 0.25 = log2(1.19)), a member of the microphthalmia (MiT/TFE) family of TFs previously shown to regulate *GBA* expression and GCase activity (33, 34). Notably, *MITF* transcription start site 2 (TSS2) (i.e. the more 3′, downstream TSS, hereafter referred to as *MITF*) and *TFEC*, two members of the MiT/TFE family, were among the 11 reproducible candidates (Fig. 2F). All those 11 candidates could also be reproduced in LN- 229^WT^ dCas9-VPR cells (Supplementary Fig. 3A). In summary, 29 out of 1,634 TFs met our primary cutoff criteria, with 11 demonstrating robust and reproducible effects on GCase activity in the lysate-based assay upon retesting. These 11 TFs were subjected to further validation.

### MITF and TFEC increase GBA transcript levels

As the enzymatic activity of GCase can be regulated at multiple levels (Supplementary Fig. 3B), we sought to differentiate between those TFs regulating GCase activity by controlling gene expression and those acting post-transcriptionally. To investigate this, we performed CRISPRa of the TFs in LN- 229^L444P^ cells and measured *GBA* transcript levels by reverse transcription quantitative PCR (RT-qPCR) and GCase protein levels through immunoblotting. We expected that most TFs would directly regulate *GBA* expression, thereby affecting GCase protein abundance. While unchanged *GBA* RNA levels make a transcriptional regulatory effect unlikely, changes in transcript levels do not necessarily imply transcriptional regulation, as alterations in RNA stability could contribute to the observed effect. For RT-qPCR, we used primers specifically targeting *GBA* to avoid amplification of the highly homologous *GBAP1* pseudogene which shares over 90% sequence identity with *GBA* (Fig. 3A).

**Fig. 3.**
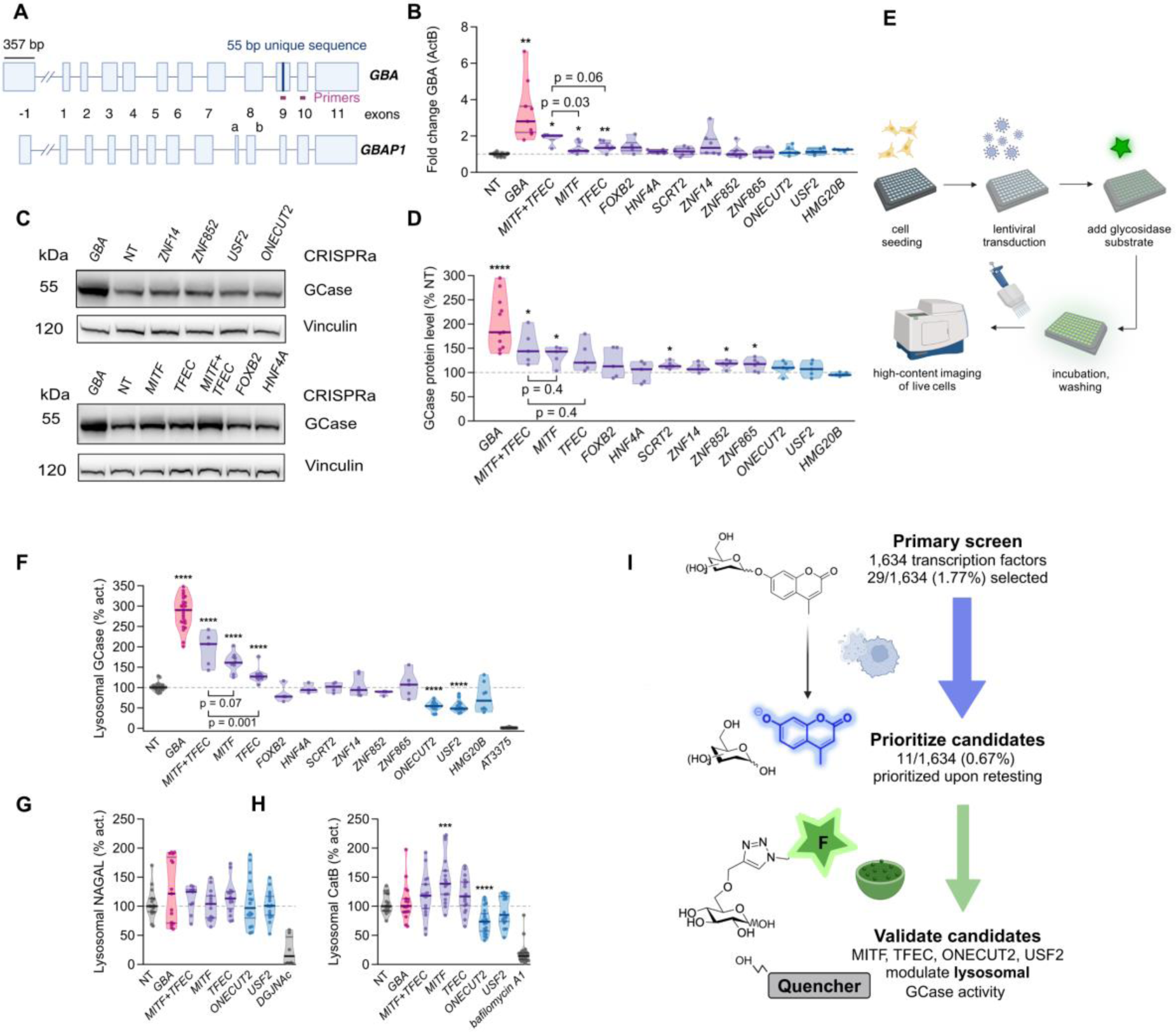
*GBA* expression, GCase protein levels, and lysosomal hydrolase activities in LN-229^L444P^ cells following activation of TFs A. *GBA* and *GBAP1* genes approximately to scale. Boxes: exons; lines: introns. Purple: 55-bp sequence unique to GBA exon 9. Pink: *GBA* RT-PCR primers (24). **B**. Fold changes *GBA* mRNA (RT-qPCR) after CRISPRa of TFs in LN- 229L444P cells. 𝛽-actin (ActB) was used for normalization (N = 3-8 repeats). **C**. GCase protein levels after CRISPRa of TFs in LN-229^L444P^ cells. Cells were harvested 5 days post-transduction. **D**. GCase protein quantification after CRIPSRa of TFs in LN-229^L444P^ cells (N = 5-11 experiments). **E**. Microscopy-based assessment of lysosomal hydrolase activities in live LN- 229L444P cells. **F**. GCase activity assessed by LysoFQ-GBA in live LN-229^L444P^ cells after CRISPRa (N = 3-9 experiments). **G**. α-N-acetylgalactosaminidase (NAGAL) activity in live LN-229^L444P^ cells after CRISPRa of TFs. DGJNAc: NAGAL inhibitor (3 experiments). H. Cathepsin B activity assessed in live LN-229^L444P^ cells after CRISPRa of TFs. Bafilomycin A1: v-ATPase inhibitor (N = 3 experiments). I. Hit selection process. Solid lines in C and D: medians, 25%, and 75% quartiles. Medians of NT controls: 100%. Unpaired t-test.

CRISPRa of *MITF* and *TFEC* significantly increased *GBA* transcript levels (fold changes 1.18 and 1.35, respectively) (Fig. 3B). As *GBAP1* was found to regulate *GBA* levels through miR-22-3p sequestration (Straniero et al., 2017), we also assessed transcript levels of *GBAP1* after CRISPRa of our candidates, which were unchanged (Supplementary Fig. 3C). At the protein level, CRISPRa of *MITF*, *ZNF865*, *ZNF852*, and *SCRT2* also increased GCase protein levels (135%, 125%, 120%, and 115% of the NT control, respectively) (Fig. 3C-D). In summary, CRISPRa of *MITF* and *TFEC* altered *GBA* transcript levels, suggesting their effects on GCase activity are at least partly mediated through the regulation of *GBA* RNA abundance. Furthermore, CRISPRa of *MITF*, *ZNF852*, *ZNF865*, and *SCRT2* increased GCase protein levels, indicating that changes in protein abundance contribute to the observed increase in GCase activity in the lysate-based assay.

### Four TFs modulate lysosomal GCase activity

While lysate-based assays report on total cellular enzyme activity, they require the disruption of the lysosomal microenvironment (27). Accordingly, we wanted to corroborate the effect of our candidate TFs on GCase activity in its native lysosomal environment. To quantify lysosomal GCase activity in live LN- 229^L444P^ cells following CRISPRa of our 11 candidates, we employed the fluorescence-quenched cell-active substrate LysoFQ-GBA, which specifically and quantitatively reports on GCase activity in lysosomes after being taken up via endocytosis (27) (Fig. 3E, Supplementary Fig. 4). The fluorescence intensities observed in wells treated with the highly selective GCase inhibitor AT3375 were <5% of the values in NT-treated control wells, confirming that the fluorescent signal originated from the turnover of LysoFQ-GBA by GCase. Among the 11 TFs, two up-regulators (*MITF* and *TFEC*) and two down-regulators (*USF2* and *ONECUT2*) significantly changed lysosomal GCase activity (Fig. 3F).

We then asked if the candidate TFs exclusively regulate GCase activity or exert a broader effect on lysosomal hydrolase activities, as was described for *TFEB* and *TFE3* (13, 34). To do this, we used the bis- acetal-based (BAB) fluorescence-quenched substrate for α-N-acetylgalactosaminidase (NAGAL-BABS) (35) and the Magic Red substrate for cathepsin B (CatB) in live LN-229^L444P^ cells. No effect on lysosomal NAGAL activity was observed following the activation of our TFs. However, activation of *MITF* increased lysosomal CatB activity whereas activation of *ONECUT2* resulted in decreased CatB activity (Fig. 3G-H); hence the directionality of these effects paralleled those seen for GCase activity. The effects of *TFEC* and *USF2* activation on CatB activity showed trends similar to their effects on GCase activity measured using the lysate assay (p = 0.076 and p = 0.1; Fig. 3H).

Overall, among the 11 modulators of GCase activity identified using the lysate-based assay, CRISPRa of *MITF*, *TFEC*, *ONECUT2*, and *USF2* significantly altered lysosomal GCase activity in live LN-229^L444P^ cells (Fig. 3I). Additionally, besides their impact on GCase, CRISPRa of *MITF* and *ONECUT2* significantly affected CatB activity, indicating a regulatory influence of these TFs on lysosomal hydrolases beyond GCase (Supplementary Fig. 5A-E). The absence of effects on NAGAL activity suggests that different lysosomal hydrolases are governed by distinct mechanisms and TFs.

### MITF, TFEC, and USF2 alter lysosomal GCase activity in iPSC-derived forebrain neurons

To determine whether the effect of the four candidate TFs on GCase activity extends to more clinically relevant cellular models, we introduced lentiviral overexpression vectors of these candidates into iPSC-derived forebrain neurons derived from healthy controls, from a patient with PD carrying the heterozygous *GBA^N370S/WT^*mutation resulting in a moderate decrease in GCase activity (around 50 % of WT) and from a GD patient with the compound heterozygous mutation *GBA^L444P/P415R^* resulting in a drastically reduced GCase activity (around 10 % of WT) (27). GCase activity was assessed in live cells using LysoFQ-GBA 7- and 14-days post-transduction. In line with the findings in LN-229^L444P^ cells following CRISPRa of candidate TFs, overexpression of the MITF-M isoform resulted in the strongest increase in lysosomal GCase activity across all three lines, followed by TFEC overexpression. Overexpression of USF2 decreased lysosomal GCase activity across all three lines (to 80% and 90% of the empty vector in the WT and PD/GD lines, respectively), while overexpression of ONECUT2 did not affect enzymatic activity, except for a trend toward a decrease in the *GBA^L444P/P415R^* neurons 14 days post-transduction (Fig. 4A-F, Supplementary Fig. 4F).

**Fig. 4.**
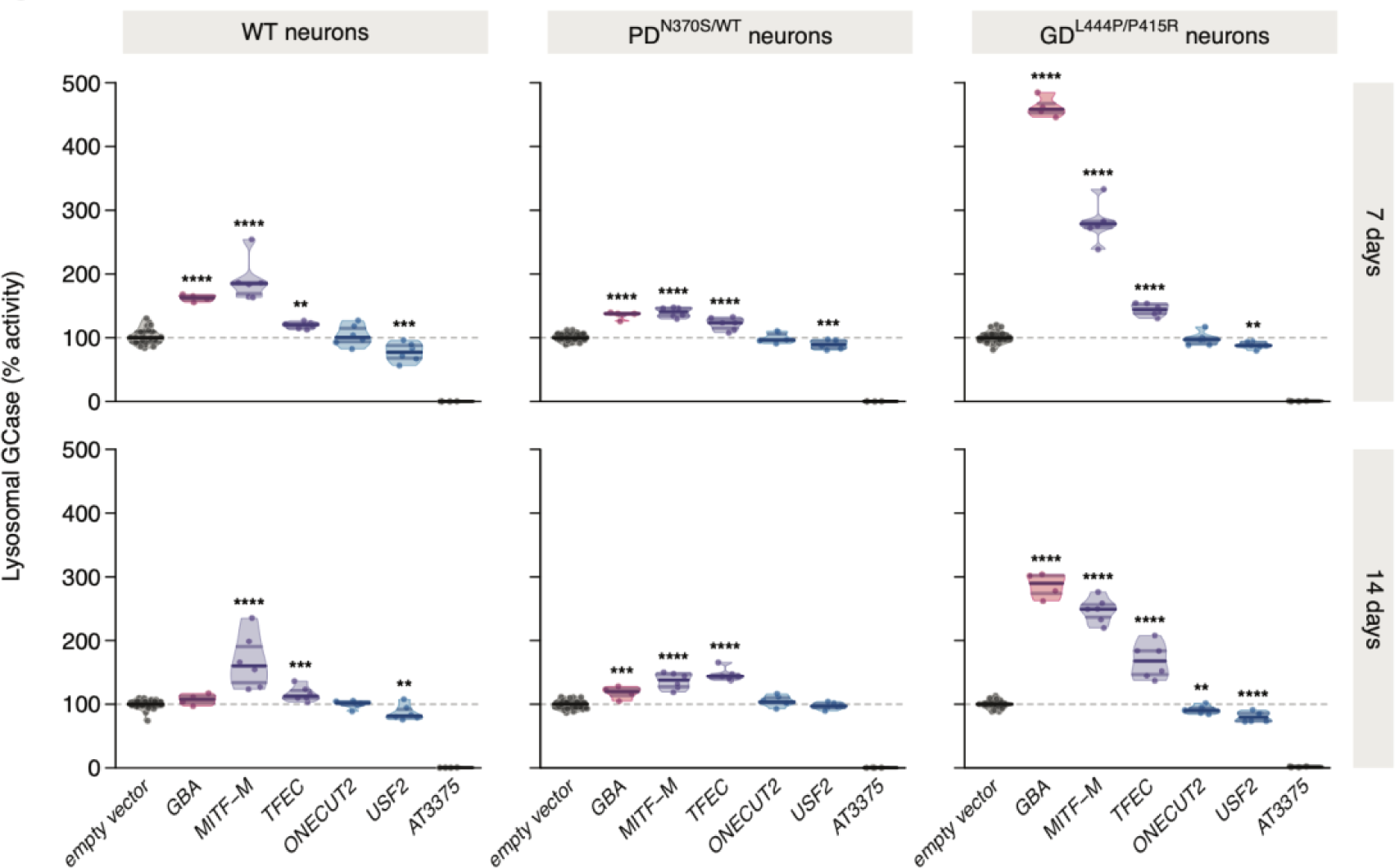
Lysosomal GCase in iPSC-derived forebrain neurons. GCase activity assessed with LysoFQ-GBA in live neurons derived from a healthy individual (WT neurons), from a Parkinson patient carrying a heterozygous *GBA* mutation (PD^N370S/WT^) or a Gaucher patient with compound heterozygous *GBA* mutations (GD^L444P/P415R^) after overexpression of candidate TFs at 7 (top row) or 14 days (bottom row) post-transduction. Six replicates/candidate and 26 replicates for the empty vector. Lines: medians and 25-75% quartiles. Medians of empty vector-infected neurons: 100%. Unpaired t-test

In summary, the effect of MITF, TFEC, and USF2 on lysosomal GCase activity was replicated in iPSC-derived neurons from both *GBA* WT and *GBA* mutant individuals harboring the two most prevalent pathogenic variants (namely, L444P and N370S), while ONECUT2 overexpression did not significantly alter lysosomal GCase activity. This discrepancy may be attributable to cell-type, isoform-and/or mutation-specific effects of ONECUT2 on enzymatic activity.

### USF2 is a bidirectional regulator of GCase abundance and activity

To assess whether the candidate genes regulate GCase activity bidirectionally, we ablated them using CRISPR in LN-229^L444P^ and LN-229^WT^ cells stably expressing Cas9. While CRISPRa of *USF2* decreased GCase activity, ablating USF2 increased GCase activity in LN-229^L444P^ in both lysate and live-cell assays (Fig. 5A and B). L444P GCase protein levels increased to 131% of the NT control following ablation of USF2 (Fig. 5C and D). The ablation efficiency was high, resulting in >75% reduction of USF2 protein levels (Fig. 5E). The effect of USF2 ablation on GCase activity and abundance was preserved in LN-229^WT^ cells (Fig. 5F and G). Notably, USF2 exhibits functional overlap with its homolog USF1, and they bind DNA either as homo- or heterodimers (36). To assess the effects of *USF1*, *USF2*, and *USF1/USF2* double-knockout, transcriptomic profiling was performed in triplicates of LN-229^L444P^ cells following CRISPR ablation. Remarkably, *GBA* emerged among the top five differentially expressed genes (DEGs) in the *USF1/USF2* double-knockout samples (Fig. 5H-I). Transcript levels of other lysosomal hydrolases (*CTSB*, *CTSD*, *GLA*) and lysosomal membrane proteins (e.g., *MCOLN1*) were also altered. Furthermore, *USF1/USF2* double-knockout also significantly increased transcript levels of *SCARB2* (encoding the GCase transport protein LIMP-2) and *PSAP (*encoding the GCase activating protein saposin C) which were previously described as modulators of GCase (37) (Fig. 5I). In addition, subunits of the lysosomal v-ATPase were differentially expressed in both *USF2* and *USF1/2* ablated samples, consistent with phagosome acidification (GO:0090383) and lysosomal membrane (GO:0005765) being among the top five gene ontology (GO) terms identified in the overrepresentation analysis (ORA) (Fig. 5I).

**Fig. 5.**
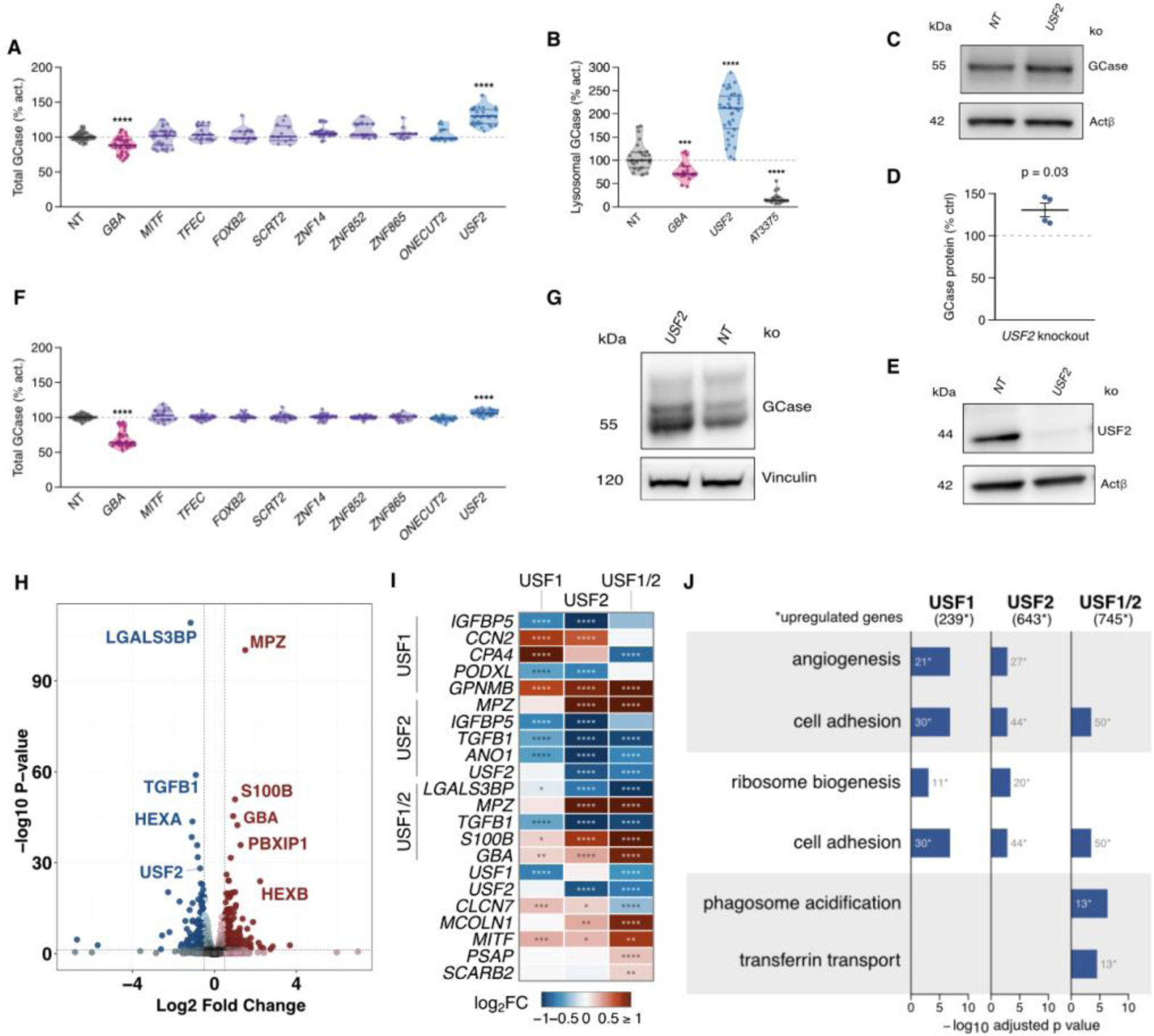
USF2 ablation increases *GBA* transcription and GCase activity/abundance. **A**. GCase activity in cell lysates of LN-229^L444P^ expressing Cas9 following ablations of TFs (N = 3 experiments). B. GCase activity in live LN-229^L444P^ cells expressing Cas9 assessed with LysoFQ-GBA after USF2 or GBA ablation (N = 3). **C**. GCase immunoblot after USF2 ablation in LN-229^L444P^ cells expressing Cas9. **D**. GCase protein levels after ablation of USF2 in LN-229^L444P^ expressing Cas9 (N = 4). **E**. Immunoblot demonstrating 75% decrease of USF2 in LN-229^L444P^ cells expressing Cas9 after ablation. **F**. GCase activity in LN-229^WT^ cell lysates after ablating candidate TFs (N = 3). **G**. Immunoblot for GCase after ablating USF2 in LN-229^WT^ expressing Cas9. **H**. Selected top DEGs following transcriptomic profiling after ablation of USF1 and USF2 in LN-229^L444P^ cells. I. Transcriptional profiling of select genes following ablation of USF1, USF2, or USF1/2 in LN-229^L444P^ cells. Horizontal axes: log2 fold-change (log2FC) in gene expression. **J**. Overrepresented gene-ontology terms. Bar length: -log10 adjusted p. (H-J: N = 3 replicates). Median values of NT controls: 100%. Solid lines: medians, 25%, and 75% quartiles

In summary, while CRISPRa of *USF2* decreased GCase activity, its ablation increased the abundance and activity of GCase in both LN-229^L444P^ and LN-229^WT^ cells, establishing *USF2* as a bi-directional regulator of GCase activity not restricted to the *GBA* L444P mutation. Furthermore, concurrent ablation of *USF1* and *USF2* enhanced transcript levels of *GBA*, other lysosomal hydrolases, and lysosomal membrane proteins, reaffirming the involvement of USFs in the modulation of lysosomal gene expression, as previously documented in murine neurons (36).

### Transcriptomic profiling suggests a role for lysosomal pH regulation by MITF and TFEC

To obtain an unbiased view of the transcriptional changes induced by TF overexpression and to identify potential shared regulatory pathways, we profiled the transcriptome of LN-229^L444P^ cells in triplicate following lentiviral transduction of the relevant CRISPRa qgRNAs. In agreement with the RT-qPCR results, *GBA* RNA levels were differentially expressed upon CRISPRa of *MITF* and *TFEC* (Supplementary Fig. 6A). CRISPRa of *MITF* increased the expression of multiple lysosomal genes, including *CTSB*, reminiscent of the effects of *TFEB* and *TFE3* (13). Additionally, CRISPRa of *MITF* resulted in the differential expression of *TFEC*, *TFE3,* and *USF2* (Fig. 6A). ORA revealed shared GO terms related to proton-transporting ATPase activity (GO:0046961), proton transmembrane transporter activity (GO:0015078), and phagosome acidification (GO:0090383) in samples in which *MITF* and *TFEC* expression was activated. DEGs in those GO terms included numerous subunits of the lysosomal v-ATPase (Fig. 6B-C). Thus, beyond the regulation of *GBA* expression, *MITF* and *TFEC* may indirectly modulate GCase activity through lysosomal pH regulation. The altered expression of *TFEC*, *TFE3*, and *USF2* upon CRISPRa of *MITF* may suggest the existence of transcriptional networks or feedback loops in which *MITF* plays a dominant role.

**Fig. 6.**
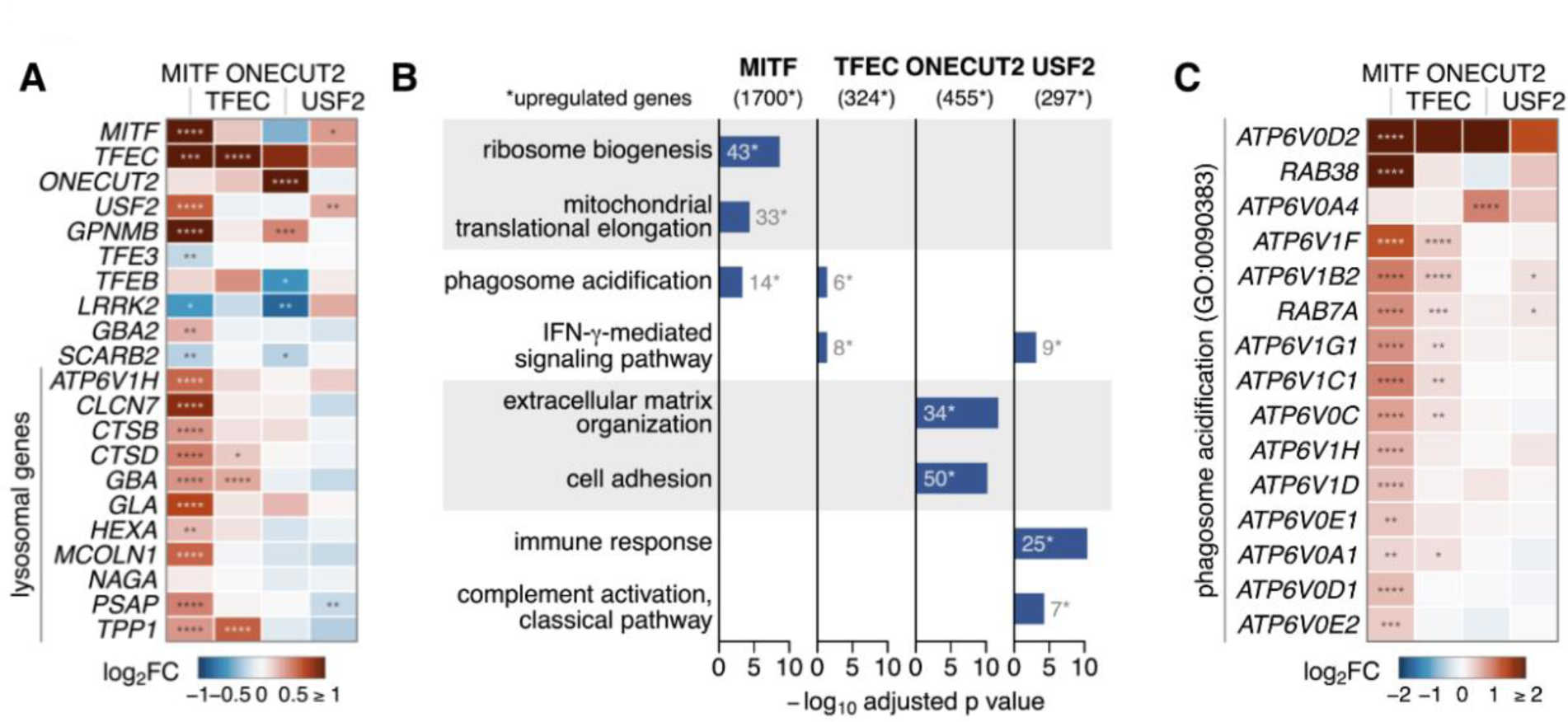
Transcriptional versus post-transcriptional mechanisms involved in *GBA* regulation. A. Transcriptional profiling of selected genes of interest following CRISPRa of candidate TFs in LN-229^L444P^ cells. Horizontal axes: log2 fold-change (log2FC) in gene expression. **B**. Top gene ontology terms of overrepresentation analysis (biological processes). Bar length:-log10 adjusted p. **C**. Differentially expressed genes of GO term 0090383 (phagosome acidification) (N = 3 replicates per

### PLEKHG4 and PLEKHG4B may mediate the downstream effects of ONECUT2 on GCase activity

RT-qPCR and RNASeq showed that *ONECUT2* activation did not alter *GBA* RNA levels (Fig. 3B, Supplementary Fig. 6A), indicating this TF regulates GCase posttranscriptionally. Many of the top DEGs identified after *ONECUT2* overexpression are predicted to be involved in vesicle trafficking (Fig. 7A). Therefore, we hypothesized that *ONECUT2* regulates GCase activity by influencing its trafficking to the lysosome or via lysosomal exocytosis. To identify potential mediators of these effects, we tested GCase activity upon activation of each of the top 30 DEGs identified through transcriptomic profiling. LN-229^L444P^ and LN-229^WT^ cells stably expressing dCas9-VPR were transduced with qgRNAs targeting the top 30 DEGs, and GCase activity was assessed five days post-transduction in cell lysates with 4MUG-Glc. Although changes in total GCase activity were modest and the overlap between LN- 229^WT^ and LN-229^L444P^ was limited, CRISPRa of *PLEKHG4* and *PLEKHG4B* showed a significant reduction of GCase activity in LN-229^WT^ cells (96 and 97% of the NT control) (Fig. 7B-C). Both genes encode guanine nucleotide exchange factors (GEFs) implicated in regulating cytoskeleton dynamics at the Golgi apparatus (38, 39). Since alterations in trafficking are likely to induce stronger changes in lysosomal rather than total GCase activity, we assessed the effect of CRISPRa of *PLEKHG4* and *PLEKHG4B* on lysosomal GCase activity using LysoFQ-GBA. Indeed, activation of *PLEKHG4* and *PLEKHG4B* significantly decreased lysosomal GCase activity in both LN-229^WT^ and LN-229^L444P^ cells (mean of 84 and 80% or 83 and 70% of the NT control, respectively) (Fig. 7D). Additionally, we evaluated alterations in protein abundance following CRISPRa of *ONECUT2* by mass spectrometry analysis in LN-229^L444P^ cells. Six of the top DEGs (RAB36, TPM1, A2M, NGFR, COL11A2, and TXNIP) exhibited changes in protein abundance (p < 0.01 with fold changes > 1.5), with RAB36 predicted to be involved in vesicle trafficking. Consistent with immunoblotting results, GCase abundance was unaffected by *ONECUT2* activation (Supplementary Fig. 7). Intriguingly, MITF protein abundance was decreased after CRISPRa of *ONECUT2* (log_2_FC = -0.74, p = 0.002)

**Fig. 7.**
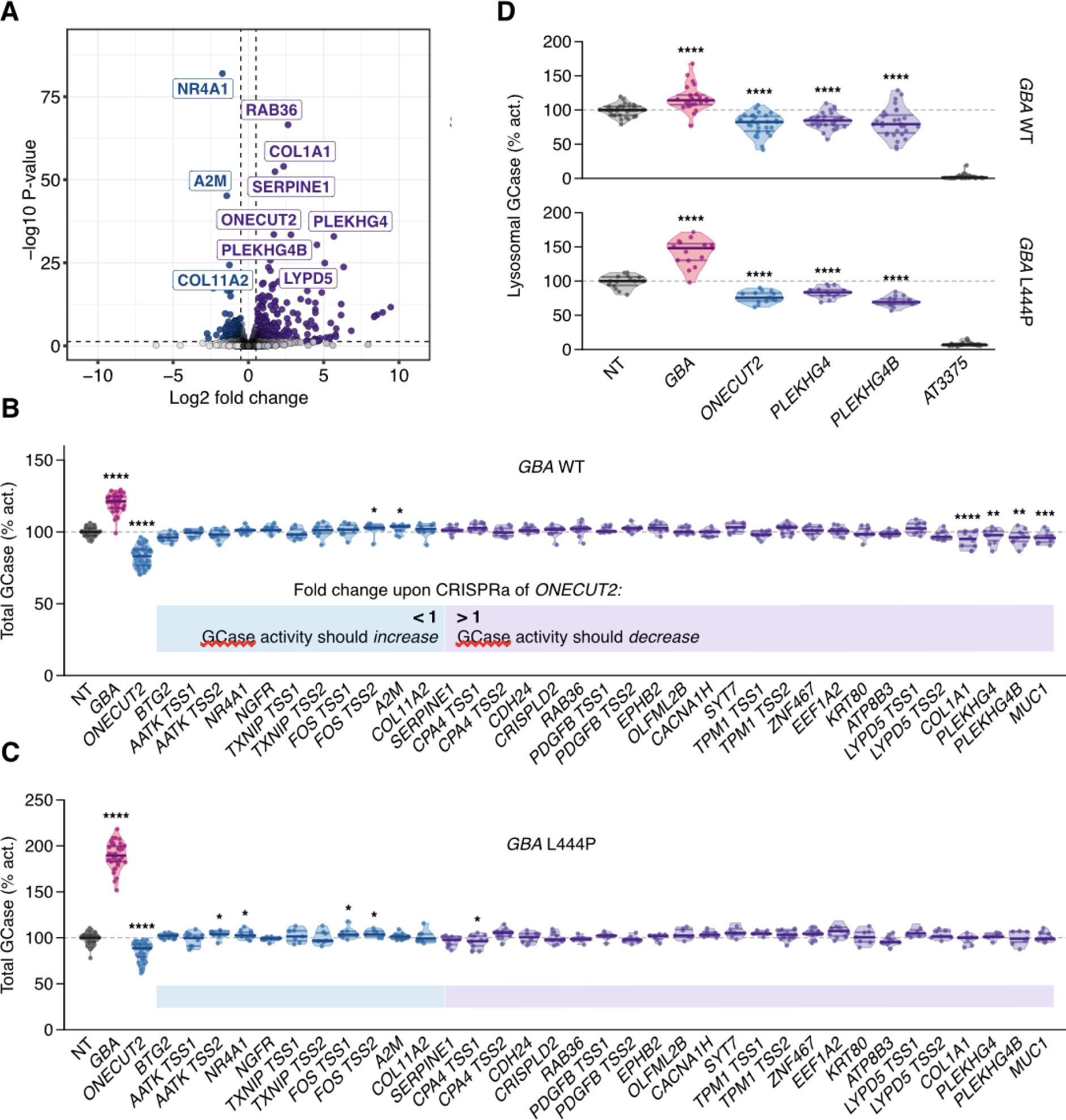
Identification of downstream mediators of ONECUT2’s effect on GCase activity. A. Transcriptomic profiling of selected DEGs following after CRISPRa of *ONECUT2* in LN-229^L444P^ cells (N = 3 replicates). **B-C**. Enzymatic activity in cell lysates following activation of top 30 DEGs in LN-229^WT^ (**B**) or LN-229^L444P^ (**C)** (N=7 to 10). **D**. Lysosomal enzymatic activity assessed with LysoFQ-GBA following CRISPRa of *PLEKHG4* or PLEKHG4B in LN-229^WT^ and LN-229^L444P^ cells (N = 3). Median values of the NT controls set as 100%. Solid lines: medians and quartiles. Unpaired t-test.

**Table 1:**
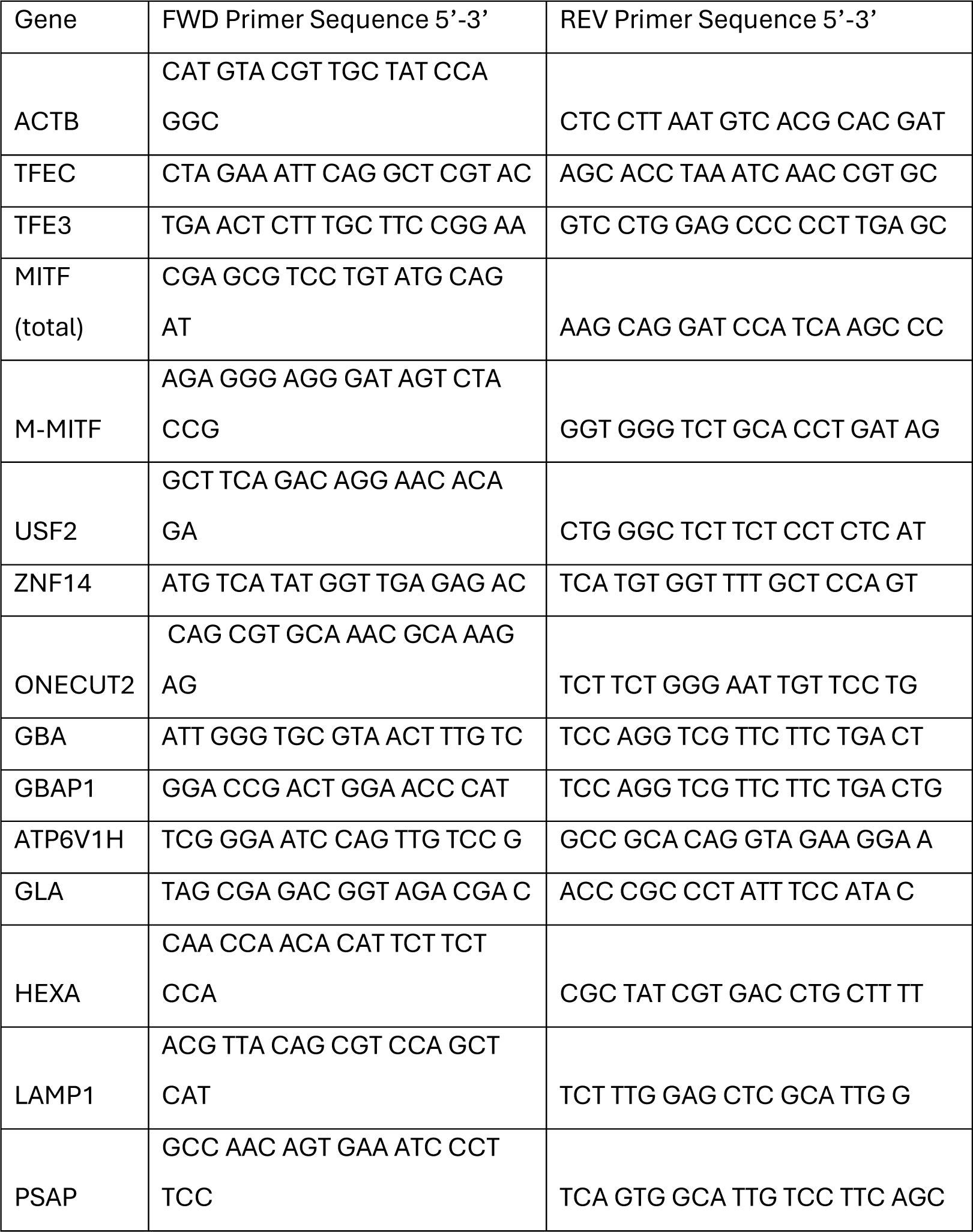
List of primers for RT-qPCR.

In summary, screening the top 30 DEGs identified upon CRISPRa of *ONECUT2* for their effect on GCase activity suggests that the GEFs PLEKHG4 and PLEKHG4B mediate, at least partially, the effect of *ONECUT2* activation on GCase activity. Further, consistent with the hypothesis that *ONECUT2* might alter trafficking of GCase to the lysosome, the effects on GCase were more pronounced when lysosomal enzymatic activity was assessed in live cells compared to assessing total GCase activity in cell lysates (Fig. 7B-D).

### Co-activation of MITF and TFEC amplifies GBA expression, GCase abundance, and activity

Since members of the MiT/TFE family bind DNA as both homo- and heterodimers (40), we explored whether simultaneous activation of *MITF* and *TFEC*, hereafter referred to as co-activation, would yield a stronger effect on *GBA* transcript levels, GCase abundance, and activity compared to individual activation. LN-229^L444P^ dCas9-VPR cells were transduced with qgRNAs targeting *TFEC* or *MITF* individually or with NT controls (MOI: 4). For co-activation, cells were transduced with qgRNAs targeting *TFEC* and *MITF* (MOI: 4 each) or with NT controls (MOI: 8). *GBA* transcripts rose more strongly upon co-activation than upon individual activation of *MITF* and *TFEC* (2x, 1.2x and 1.3x, respectively) as measured by RT-qPCR and bulk RNASeq (Fig. 3B, Supplementary Fig. 6B). Moreover, co-activation of *MITF* and *TFEC* led to the most pronounced increase in GCase protein levels (mean increase of 150% for co-activation versus 130% for individual activations compared to the NT control) (Figures 3D). Consistent with their effects on *GBA* transcript and GCase protein levels, co-activation of *MITF* and *TFEC* more substantially increased lysosomal GCase activity compared to their individual activation (173% vs. 157% and 147% of the NT control, respectively). However, the difference between co- and individual activation on lysosomal GCase activity was only significant for TFEC (p=0.001) (Fig. 3F). To summarize, co-activation of *MITF* and *TFEC* resulted in a synergistic effect on *GBA* expression, GCase abundance and activity, with the most substantial increase detected at the RNA level.

## Discussion

We used unbiased forward genetics to systematically define TFs modifying GCase activity in the presence of the pathogenic *GBA* L444P variant. Currently available CRISPRa libraries, unlike T.gonfio, are produced in a pooled format which limits their utility to assays reliant on selectable phenotypes like cell death, cell survival, or expression of cell-surface markers, and renders them unsuitable for complex biochemical screens as the one presented here. Therefore, we screened all human transcription factors by employing a subset of the genome-wide T.gonfio arrayed CRISPRa library (18).

In principle, overexpression of cellular genes can be achieved by transducing the genes of interest under transcriptional control of strong heterologous promoters. However, CRISPRa offers several advantages over traditional overexpression strategies in that it allows for the upregulation of TFs within their native context, thus preserving the cellular post-transcriptional processing. We had considered assaying GCase upon CRISPR-mediated arrayed gene ablation instead of gene activation. However, disrupting single genes one by one is not effective in identifying redundant pathways, as the ablation of one gene may not suffice to induce a detectable phenotype. Moreover, ablating genes that are not endogenously expressed within the cell line under study cannot induce a phenotype. Lastly, unlike CRISPR ablation, CRISPRa enables studying the effect of essential genes on pathways of interest without inducing cell death, thereby allowing interrogation of a larger number of genes.

Since transcriptional regulation has been predicted to influence GCase protein abundance and GCase function is associated with disease, we elected to screen for enzymatic activity in cell lysates. While lysate-based assays yield robust and reproducible results, they lack topological resolution since they report on total cellular GCase. Hence, these assays may overestimate residual enzymatic activity, especially in the context of certain *GBA* mutations like the L444P variant, as the misfolded enzyme may be retained in the endoplasmic reticulum before being targeted to the proteasome. Moreover, changes in the lysosomal microenvironment may go undetected as lysosomes get disrupted (Supplementary Fig. 5C). Hence, TFs identified in the primary screen were validated in live cells by using the cell-active substrate LysoFQ-GBA to measure the activity of the physiologically relevant lysosomal GCase fraction (27). Employing these approaches, we validated *MITF*, *TFEC*, *USF2*, and *ONECUT2* as regulators of lysosomal GCase activity.

### The bHLH-LZ family members MITF, TFEC, and USF2 regulate GBA transcript levels

The identification of MITF and TFEC, members of the MiT/TFE family which includes TFEB and TFE3, is not unexpected and lends confidence to our findings. In contrast, USF2 and ONECUT2 have not been previously linked to the regulation of GCase activity. Except for ONECUT2, these are all members of the basic helix-loop-helix leucine zipper (bHLH-LZ) family of TFs, which recognize DNA sequences known as enhancer (E)-boxes comprised of a CANNTG sequence. The CLEAR element harbors an E-box (13), and MITF, TFEC, and USF2 are all predicted to bind to the CLEAR motif according to the JASPAR database of TF binding profiles (41). In line with this prediction, CRISPRa of *MITF* and *TFEC* and ablation for USF2 changed *GBA* transcript levels as assessed by RT-qPCR or bulk RNASeq.

Transcriptomic profiling revealed differential expression of lysosomal genes other than *GBA* upon *MITF* or *TFEC* activation and USF2 ablation. CRISPRa of *MITF* also increased lysosomal cathepsin B activity. Besides regulation of *GBA* and *CTSB* transcript levels, transcriptomic profiling suggested that further factors, such as the regulation of lysosomal pH, might contribute to the increase in lysosomal hydrolase activities upon CRISPRa of *MITF*. However, this effect would have been missed in our primary screen employing a lysate-based assay, as lysosomes are disrupted, and the artificial substrate is provided in an acidic buffer. Hence, more TFs affecting lysosomal pH may exist beyond MITF.

Co-activation of *MITF* and *TFEC* led to a more pronounced increase in *GBA* RNA, GCase protein, and activity levels as compared to their individual activation, suggesting that heterodimer formation of MITF/TFEC results in higher affinity binding to the *GBA* promoter and/or more pronounced activation of *GBA* expression compared to their binding as homodimers. It remains to be seen in which cell type heterodimer formation is functionally relevant. Nevertheless, the simultaneous activation of several genes represents a useful approach to assessing the additive or synergistic effects of genes predicted to be involved in the same pathways.

We identified *USF2* as a bidirectional regulator of GCase activity in both LN-229^L444P^ and LN-229^WT^ cells. The upstream factor USF2 and its homolog USF1 are evolutionarily conserved, ubiquitously expressed, and bind to DNA as homo- or heterodimers. USFs are implicated in the regulation of various cellular processes including proliferation, immune response, lipid metabolism, and histone acetylation (42), and USFs have been proposed as regulators of lysosomal function in murine neurons (36). As members of the bHLH-LZ family, USFs may compete with the MiT/TFE family members for binding to the CLEAR motifs in the promoters of lysosomal genes, thereby antagonizing their effects. A similar mechanism has been proposed for MYC, another member of the bHLH-LZ family (36, 43). Notably, a single nucleotide variant in IL-10 which may alter the USF2 binding site was proposed to explain phenotypic variability in Fabry Disease, another lysosomal storage disorder (44).

CRISPRa of all other identified TFs did not alter *GBA* transcript levels, and no predicted binding sites were found in the *GBA* promoter according to the JASPAR database. These intriguing results suggest that GCase is regulated through post-transcriptional mechanisms, some of which can be targeted by the TFs identified by our screen. A special case is represented by the zinc finger proteins *ZNF852*, *ZNF865*, and *SCRT2* which increased GCase protein levels although *GBA* RNA levels and lysosomal GCase activity measured with LysoFQ-GBA remained unchanged. This could point to alterations in GCase protein stability, as the mutant enzyme might partially evade premature proteasomal degradation but fail to enter the lysosome due to a trafficking defect.

### ONECUT2 is a posttranscriptional regulator of GCase activity

The *ONECUT* family members (ONECUT1–3) participate in the development of the liver, pancreas, and retina but are not associated with lysosomal function (45, 46). Although ONECUT2 binds to the MITF promoter in melanocytes (47), *GBA* RNA and GCase protein levels remained unchanged upon CRISPRa of *ONECUT2*, suggesting that it does not regulate *GBA* transcription. In addition to it decreasing GCase activity, CRISPRa of *ONECUT2* also decreased lysosomal CatB activity.

Transcriptomic profiling also did not show a significant decrease in *CTSB* expression, pointing to a broader regulatory effect on lysosomal function. Supporting this idea, several genes associated with vesicle trafficking were among the top DEGs (Fig. 7A). Thus, *ONECUT2* may regulate GCase activity by influencing its trafficking. Despite small effect sizes and the likelihood of trafficking defects being overlooked in a lysate-based assay, screening of the top 30 DEGs identified *PLEKHG4* and *PLEKHG4B* as possible downstream mediators. Indeed, CRISPRa of *PLEKHG4*, *PLEKHG4B,* and *PLEKHG4+PLEKHG4B* demonstrated a stronger effect on lysosomal GCase activity (assessed with LysoFQ-GBA) than on total cellular GCase activity (assessed with 4MU-Glc) in both LN-229^WT^ and LN- 229^L444P^ cells. Since *PLEKHG4* and *PLEKHG4B* are guanine nucleotide exchange factors (GEFs) implicated in cytoskeleton dynamics at the Golgi apparatus, their overexpression may enhance GCase trafficking and lysosomal GCase activity (38, 39).

### Limitations of this study

The LN-229 cancer cell line was primarily chosen for practical reasons. LN-229 cells are polyploid and exhibit genomic instability due to a *TP53* mutation, which may affect the generalizability of our results. While the effects of MITF, TFEC, and USF2 overexpression onto lysosomal GCase activity were confirmed in iPSC-derived forebrain neurons carrying the highly prevalent N370S and L444P *GBA* variants, ONECUT2 was ineffective. Possible reasons might be related to the experimental design. Overexpression vectors, unlike CRISPRa, only increase the expression of specific isoforms that may not be relevant to the cell type being tested. Alternatively, the ONECUT2 effectors might be cell-type specific and absent from glutamatergic neurons, or the phenotype might be mutation-specific. As the tested GD-derived neurons harboring the *GBA*^L444P/P415R^ mutation exhibit low baseline GCase activity, any further decrease induced by ONECUT2 might be below the detection limit of our assay.

It remains to be seen if these effects are of physiological relevance, as both endogenous MITF and TFEC levels are predicted to be low in neurons. Given the emerging role of neuroinflammation in the pathogenesis of neuronopathic GD and PD, testing these candidates in microglia will be of interest (48, 49). We introduced the pathogenic *GBA* L444P mutation into LN-229 cells because it is the most prevalent pan-ethnic point mutation associated with severe nervous system involvement unamenable to enzyme replacement therapy in GD patients (6). Due to their low baseline enzymatic activity, we anticipated a more pronounced increase in GCase activity in LN-229^L444P^ upon perturbation of regulatory pathways, which we expected would enhance the dynamic range of the lysate assay.

However, changes in GCase activity may remain below detection limits due to the continued extensive proteasomal degradation associated with this mutation. Moreover, the >300 distinct *GBA* mutations that impair its enzymatic activity do so through mechanisms extending beyond proteasomal degradation, including disrupted interaction with the activating protein saponin C. Thus, the effect of the identified TFs on GCase activity may vary depending on the specific mutation (50).

Curiously, TFEB and TFE3, two TFs known to regulate the expression of numerous lysosomal genes including *GBA* (34), were not among our top candidates. Constitutive activation of the mTORC1 pathway in glioblastoma cells may explain their absence, as phosphorylation by mTORC1 prevents their nuclear translocation (51). In contrast, TFEC and MITF-M (the isoform corresponding to CRISPRa of *MITF* TSS2) lack an mTORC1 phosphorylation site and may therefore be able to enter the nucleus under such conditions (52). Moreover, functional redundancy of the MiT/TFE family members contributes to their cell-type-dependent regulation of metabolic pathways (33).

To more clearly address the functional effects of altering the expression of the TFs we identify as regulators of lysosomal GCase activity, TFs could be tested for their effect on 𝛼-synuclein aggregation, glucosyl sphingosine levels, or endoplasmic reticulum stress, which have all been associated with PD risk in the context of *GBA* mutations (2).

In conclusion, by conducting an arrayed CRISPRa screen in cell lysates, we identified four TFs that modulate lysosomal GCase activity when assessed with the cell-active substrate LysoFQ-GBA within live cells. These results point to an intricate network of TFs beyond TFEB and TFE3 that regulates lysosomal GCase activity, with mechanisms beyond direct transcriptional regulation of *GBA* being at play. The identification of *MITF*, *TFEC*, *USF2*, and *ONECUT2* as regulators of GCase activity illuminates an underappreciated interplay between transcriptional regulation of lysosomal function and PD risk. While the pharmacological modulation of transcription factors is considered challenging, the surprising discovery of an ONECUT2-mediated non-transcriptional mechanism of GCase regulation may yield targets that are more amenable to pharmacotherapy.

## Supplementary materials

### Contributions (in alphabetical order)

Aguzzi, Adriano: proposed the project, supervised its execution, secured funding, helped writing and correcting the manuscript. Armani, Andrea: supervision, experimental design, writing of manuscript.

Böck, Desirée: supervision of adenine base editing; next-generation sequencing. Chen, Xi: assistance with cell culture work; experimental design and optimization of live cell assays; writing of manuscript. Deen, Matthew: design and synthesis of LysoFQ-GBA; writing of manuscript. Dhingra, Ashutosh: experimental design and generation of lentiviruses encoding the qgRNA plasmids targeting the human TFs used for the primary screen. Frick, Lukas: design of CRISPR library; supervision and experimental design, data analysis and visualization; writing of manuscript. Gilormini, Pierre-André: supervision, experimental design of lysate-based and live cell enzyme activity assays. Ging, Kathi: generated cellular model system, designed and performed, or contributed to all experiments; writing of manuscript. Marques, Ana: assistance in cell culture, live cell assay, immunoblotting, and RT-qPCR; writing of manuscript. Pisano, Claudio: experimental design; assistance in cell culture, lysate-based assay, immunoblotting, and RT-qPCR Schlachetzki, Johannes: supervision, experimental design, writing of manuscript. Sellitto, Stefano: experimental design; assistance in cell culture; writing of manuscript. Trevisan, Chiara: assistance in primary CRISPR activation screen; assistance in cell culture; writing of manuscript. Yin, Jiang-An: design and production of CRISPR library; supervision and experimental design; writing of manuscript. Zhu, Yanping: experimental design. Heutink, Peter; Vocadlo, David; Glass, Christopher K: supervision of parts of the project, secured funding, manuscript writing

## Supporting information

Hit_list_primary_screen

## Acknowledgments

We thank Prof. em. Konrad Sandhoff for his valuable input and explanations. We thank Buket Züllig for their assistance in cell culture, immunoblotting, and RT-qPCR. We thank Lennart Opitz and Dr.

Timothy Sykes and the Functional Genomics Center Zurich for their help with the RNA-Sequencing data analysis. We thank Joana Delgado Martinez and the Centre for Microscopy and Image Analysis of the University of Zurich for her help with and access to the microscopes. Figures 1A, 2A, 3A/E/I, Supplementary Fig. 3B and 6C were created with BioRender. We thank Amicus Therapeutics for providing AT3375.

## Materials and Methods

Cell culturing

### Generation of the LN-229^L444P^ cell line

Cells were cultured in DMEM (Gibco) supplemented with 10% Fetal Bovine Serum (FBS). All experiments were performed in cells under passage 40. The *GBA* L444P mutation was introduced into the LN-229 human glioblastoma cells (CRL-2611, ATCC).

gRNA was ordered as forward and reverse primers from Microsynth (Balgach, Switzerland).

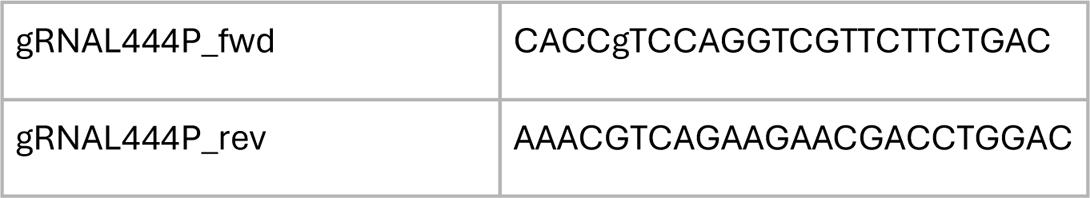

### gRNA cloning

The lentiGuide-Puro plasmid (Addgene #52963) was digested with Esp3I (BsmBI) (ER0451, Thermo Scientific).

|  |  |
| --- | --- |
| 5 µg plasmid DNA | x µL |
| Restriction enzyme (Esp3I) | 5 µL |
| 10x Tango buffer | 10 µL |
| DTT 20 mM | 5 µL |
| H<sub>2</sub>O | to 100 µL |

The reaction mix was incubated overnight at 37°C. On the next day, 0.5 µL of calf intestinal alkaline phosphatase (CIP) (M0290, New England Biolabs (NEB)) was added and the reaction was incubated for 1 h at 37°C.

The digested plasmid was gel purified (NucleoSpin Gel and PCR Clean-up, 740609.50, Macherey-Nagel).

To generate the gRNA, the forward and reverse primers were phosphorylated and annealed:

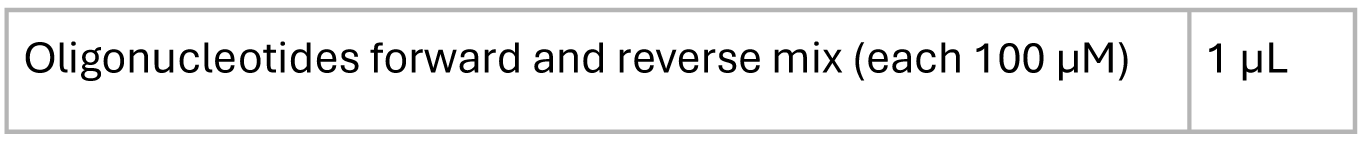
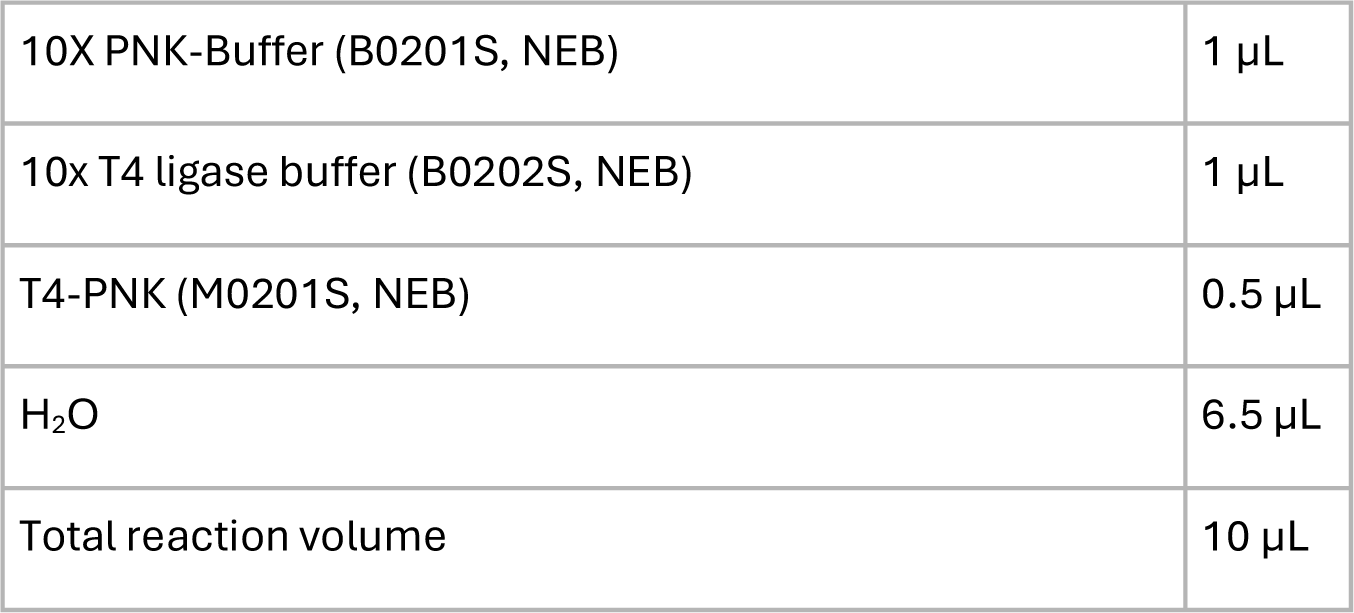

|  |  |
| --- | --- |
| Oligonucleotides forward and reverse mix (each 100 µM) | 1 µL |
| 10X PNK-Buffer (B0201S, NEB) | 1 μL |
| 10x T4 ligase buffer (B0202S, NEB) | 1 μL |
| T4-PNK (M0201S, NEB) | 0.5 μL |
| H<sub>2</sub>O | 6.5 μL |
| Total reaction volume | 10 μL |

The phosphorylation/annealing reaction was incubated for 30 min at 37°C followed by 5 min denaturation step at 95°C before ramping down the reaction to 25°C at 5°C/min.

The annealed oligonucleotides were diluted at 1:200 in sterile water. Ligation of digested plasmid and annealed oligonucleotides was performed:

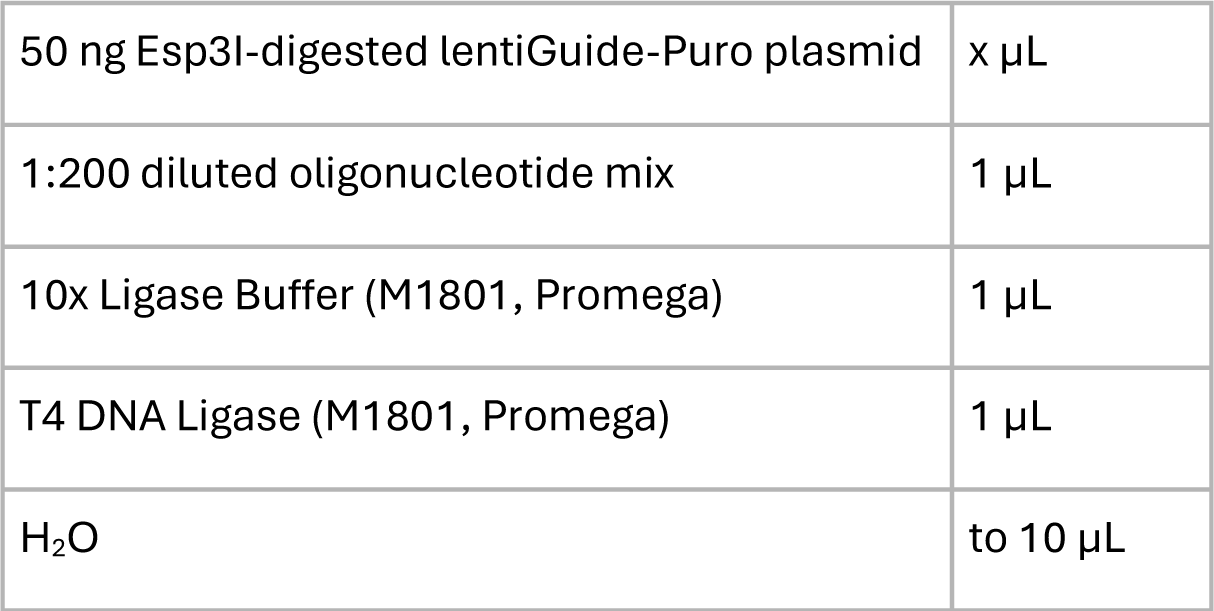

The reaction was incubated for 3 h at room temperature.

Transformation into NEB^®^ Stable Competent E. coli (C3040H) was performed as follows: 5 µL of ligation reaction were mixed with 50 µL of E. coli. The mix was kept on ice for 30 min before performing a heat shock (30 s at 42°C). After another 5 min on ice, 300 µL of S.O.C. medium (15544034, ThermoFisher) was added and E. coli were shaken at 37°C for 1 h before plating them on an ampicillin agarose plate (100 μg/mL) and incubating them overnight at 37°C. Single colonies were picked the next day and grown in LB medium containing ampicillin (100 μg/mL) for one night at 37°C in a shaker. Plasmid DNA was extracted (QIAprep Spin Miniprep Kit, 27106X4 Qiagen) and sent for Sanger sequencing (Microsynth).

5′ sequencing primer: EF-1a-F (TCAAGCCTCAGACAGTGGTTC);

3′ sequencing primer: WPRE-R (CATAGCGTAAAAGGAGCAACA).

A negative control ligation and transformation (digested vector with water in place of oligonucleotides) was run in parallel.

For base editing, LN-229 (passage 9) were transfected in 24-well plates. The adenine base editor ABE8e (Addgene #138489) (20) and the lentiGuide-Puro vector encoding gRNAL444P were co-

transfected using Lipofectamine 3000® (Thermo Fisher) in a ratio of 3:1 (i.e., 375 ng and 125 ng). Selection with Puromycin (Gibco) 1 µg/µL was started after 24 hours. Medium was changed daily. On day 5, cells from a single well were dissociated with Trypsin and diluted in 1000 µL medium. 2-50 µL of the cell suspension was transferred to 10 cm dishes to obtain single colonies. Medium containing Puromycin was changed twice weekly and single colonies were picked manually after 2-3 weeks when reaching an adequate size and expanded. Genomic DNA was extracted (DNeasy Blood & Tissue Kit, Qiagen), amplified with primers L444P_fwd and L444P_rev and sent for Sanger sequencing (Microsynth) with primers L444P_fwd2 and L444P_rev specific for *GBA* (i.e., not amplifying *GBAP1*):

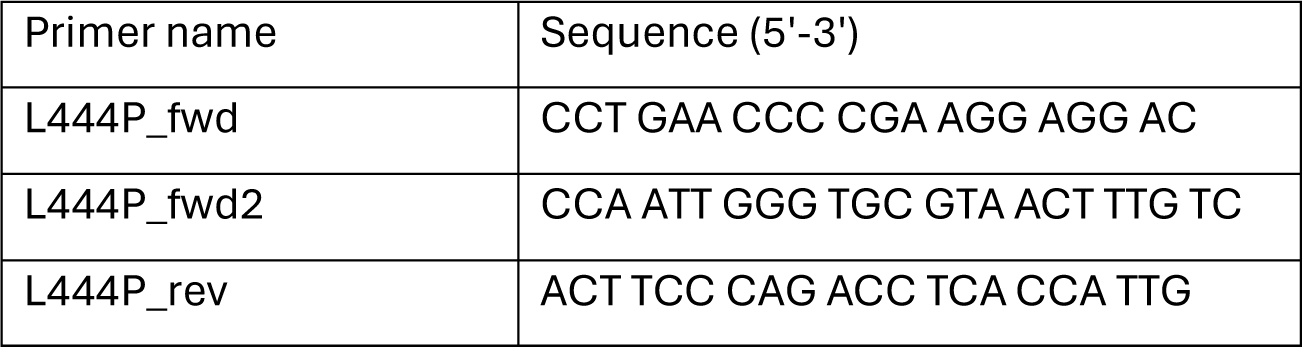

To ensure that all the alleles of the polyploid LN-229 clones were edited, Illumina® next-generation sequencing (NGS) was performed. Genomic DNA extracted from base-edited clones showing the desired point mutation with Sanger sequencing underwent PCR with primers L444P_fwd and L444P_rev to ensure that only *GBA* and not *GBAP1* was amplified. The amplicon was used as a template for the second round of PCR with primers introducing adaptors for Illumina sequencing (NGS_L444P_fwd and NGS_L444P_rev). Q5 High-Fidelity DNA Polymerase (M0491S, NEB) was used with 200 ng of template DNA in a total reaction volume of 20 µL.

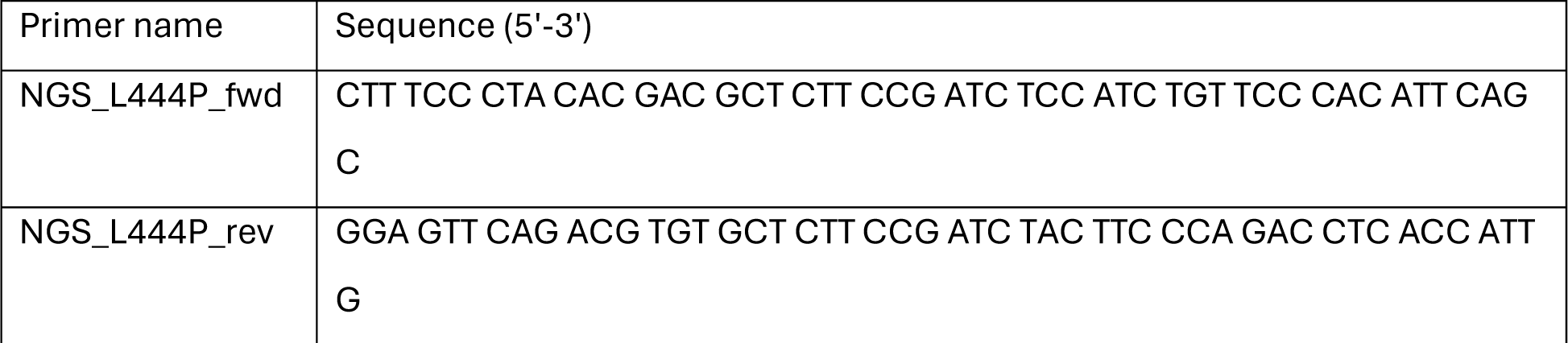

PCR products were purified using AMPure XP beads (Beckman Coulter) and subsequently amplified for eight cycles using barcoding primers. Approximately equal amounts of PCR products from each sample were pooled, gel purified (QIAquick Gel Extraction Kit, Qiagen), and quantified using a Qubit

3.0 fluorometer and the dsDNA HS Assay Kit (Thermo Fisher). Paired-end sequencing of purified libraries was performed on an Illumina® MiSeq.

Sequencing reads were demultiplexed using MiSeq Reporter (Illumina). Amplicon sequences were aligned to their reference sequences using CRISPResso2 and the tool was used to generate Figure 1B

1. (53) (Amplicon: CCATCTGTTCCCACATTCAGCAAGTTCATTCCTGAGGGCTCCCAGAGAGTGGGGCTGGTTGCCAGTCAGA

AGAACGACCTGGACGCAGTGGCACTGATGCATCCCGATGGCTCTGCTGTTGTGGTCGTGCTAAACCGGTGAGGGCAATGGTGAGGTCTGGGAAGT; sgRNA: GTCCAGGTCGTTCTTCTGAC).

### Generation of LN-229^ΔGBA^ cells

LN-229 cells were co-transfected with the plasmid encoding the sgRNAs as outlined below and hCas9 (Addgene #41815) using Lipofectamine 3000 in a ratio of 3:1. Single colonies were selected as described above. Knockout was confirmed by performing Sanger sequencing, immunoblotting, and a lysate-based enzyme activity assay.

The sequence of the sgRNAs to target the catalytic domain of *GBA* was*: The sgRNA1 (sgRNA1 sequence) oligonucleotide sequence is: 5’-* ACAGAAGTTCCAGAAAGTGA*-3’; sgRNA2 (sgRNA2 sequence, N20sg2); 5’-*GAGAGCAGCAGCATCTGTCA-3’; *sgRNA3 5’-* AGTGATGGAGCAGATACTCA *-3’; sgRNA4 5’-* GATGGAGCAGATACTCAAGG *-3’*.

### Lentiviral packaging

HEK293T cells were grown to 80-90% confluency in DMEM + 10% FBS on poly-D-lysine coated 24-well plates and transfected with the three different plasmids (Transfer plasmid, psPAX2 (#12260), VSV-G (#8454); ratios: 5:3:2) with Lipofectamine 3000 for lentivirus production. After six hours or overnight incubation, the medium was changed to virus harvesting medium (DMEM + 10% FBS + 1% BSA (Cytiva HyClone)). The supernatant containing the lentiviral particles was harvested 48-72 hours after the change to virus harvesting medium. Suspended cells or cellular debris were pelleted with centrifugation at 200 g for 5 min. Clear supernatant was titrated by FACS and stored at -80 °C in single-use aliquots.

For the titration of the lentiviral particles, HEK293T cells were grown in 24-well plates and infected with 4 mL (V) of the above-mentioned viral supernatant. Cells of two representative wells were counted at the time of infection (N). 72 hours after infection, the cells were harvested and analyzed by flow cytometry to determine the fraction of infected cells (BFP positive) and determine the viral titer (T) according to the following formula: T = (P * N)/V.

Flow cytometry analysis was performed with a BD Canto II or LSRFortessaTM Cell Analyzer at the core facility center of the University of Zurich. Data were analyzed with FlowJo.

### Generation of cell lines stably expressing dCas9-VPR or Cas9

Lentiviruses of plasmids pXPR_120 (Addgene #96917) and lentiCas9-Blast (Addgene #52962) were produced. Cells in 24-well plates at 70% confluency were infected and put under Blasticidin selection (15 µg/mL) 24 hours post-infection. Blasticidin-containing medium was changed every 48 hours and cells were gradually expanded when confluent. Successful expression of the Cas9 construct was determined by means of RT-qPCR and Western Blot of two target genes.

### Workflow of Primary Arrayed CRISPR Screen

LN-229^L444P^ cells stably expressing dCas9-VPR were cultured in T75 tissue culture flasks (TPP) in DMEM (Gibco) supplemented with 10% FBS (Takara) and 15 mg/mL Blasticidin (Gibco). In preparation for the primary screen, cells were expanded, washed with PBS (Gibco), harvested using Trypsin-EDTA 0.025% (Gibco), resuspended in DMEM supplemented as outlined above, pooled, and counted using a TC20 (BioRad) Cell Counter with trypan blue (Gibco). For the primary screen, all LN-229^L444P^ cells were at the same passage number.

2,000 LN-229^L444P^ dCas9-VPR cells were dispensed in 50 µL/well of medium using a multi-drop dispenser (MultiFlo FX, Biotek) in 384-well plates (#781091, Greiner). Plates were centrifuged at 10 g for 30 seconds (5804R, Eppendorf) and incubated overnight in a rotating tower incubator (StoreX STX, LiCONiC). Cells were transduced after 18-20 hours using a handheld electronic multichannel pipetting system (ViaFlo, Integra) with an MOI of four with lentiviruses encoding the qgRNA plasmids targeting each human TF (54). Outermost wells were spared to avoid edge effects related to evaporation. 14 wells/plate were transduced with lentiviruses encoding negative and positive controls (i.e., 14 NT ctrls and 14 *GBA* TSS1-targeting 4sgRNA). Each 4sgRNA plasmid targeting a TF was transduced in triplicates on three separate plates in the same well position. Plates were incubated in a rotating tower incubator for five days. Subsequently, one replicate plate was used to determine cell viability by applying CellTiter-Glo® (Promega) following the instructions of the manufacturer. All solutions were dispensed with a multi-drop dispenser. Briefly, medium was removed by inverting the plates and replaced with 25 µL of fresh medium and 25 µL of CellTiter-Glo® solution, both solutions were at RT. Plates were incubated on a plate shaker (ThermoMixer Comfort, Eppendorf) for two min (400 rpm shaking) at RT. After further 10 min of incubation at RT, luminescence was determined with a fluorescence plate reader (EnVision, Perkin Elmer).

GCase activity was assessed on the two remaining plates. Medium was removed by inverting the plates, and cells were lysed in 10 μL lysis buffer (McIlvaine buffer with 0.4% Triton-X), supplemented with EDTA-free cOmplete protease inhibitor. Following lysis, assay plates were incubated on a plate shaker for 10 min at 4 ° (400 rpm shaking) prior to centrifugation at 200 g for one min and incubation at 4 ° for one hour. Following incubation, plates were centrifuged under the same conditions mentioned above and 10 μL of assay buffer containing 4MUG was added. After centrifugation, plates were incubated for 1.5 hours at 37°C in a rotating tower incubator. After centrifugation, 20 μL of stop solution was added to each well. Following a final centrifugation step (300 g for 10 min), plates were read at a VersaMax Microplate Reader (Molecular Devices) (Excitation: 365 nm, Emission: 440 nm).

### Data analysis

Screening data were analyzed using an in-house developed, open-source, R-based HTS analysis pipeline developed by Dr. Lukas Frick (all documentation and codes available under: <u>github.com/Ginka21/CRISPRa_TF</u>). Initially, Z’-Factor and strictly standardized mean difference (SSMD) scores (31, 55) were calculated to assess the discrimination between the positive and NT controls. Subsequently, heat maps of individual plates were generated to examine gradients or dispensing errors. Additionally, fluorescence values were plotted to check for row and/or column effects. Correlation between duplicates was assessed by calculating Pearson’s correlation coefficient. Candidate genes were selected with the following cut-off criteria: Fold change in fluorescence intensity >1.25 as compared to NT controls and a p (t-test) of <0.05. Volcano plots and dual flashlight plots were generated for data visualization. Detailed formulas are provided in the Supplementary.

Some graphs were generated with Prism GraphPad 8.0.0.

### Reproduction of results with a higher number of replicates

The same protocol as for the primary screening was carried out. The only difference was that lentiviruses were pipetted manually.

To test for bidirectional regulation (CRISPRko of candidates), 500 cells/well stably expressing Cas9 were seeded in 384-well plates on day 0 and lysis was performed on day seven post-transduction.

### RNA extraction and Real-time quantitative PCR

Total RNA was isolated using the High Pure RNA Isolation Kit (Roche). 800-1,000 ng of RNA were reversed transcribed into cDNA using the QuantiTect Reverse Transcription Kit (Qiagen). Real-time quantitative PCR was performed with SYBR green (Roche) according to the manual with the primer sets for each gene listed in table 6. ACTB was used as the housekeeping controls. A ViiA7 Real-Time PCR system (Applied Biosystems) was used for fluorescence detection.

### Immunoblotting and deglycosylation

On day zero, 40,000 cells were seeded in 6-well plates. On day one, cells were infected with lentiviruses (MOI of 4). Medium was changed after 24 hours. Three days (CRISPRa) or four days (CRISPRko) post-transduction.

On day five (CRISPRa) or seven (CRISPRko) post-transduction, cells were lysed in RIPA buffer supplemented with Protease Inhibitor. Samples were centrifuged at 17,000x g for 10 minutes at 4°C and the supernatant was subjected to further downstream analyses. BCA assay (Pierce, Thermo Fisher) was used to determine the total protein concentration of each sample. For immunoblotting, samples were diluted in RIPA buffer to obtain the same total protein concentration. Deglycosylation was performed with either PNGase F or EndoH (both from NEB), following the protocol of the manufacturer.

Samples were boiled at 100°C for 10 min after addition of NuPAGE 4x LDS loading buffer (Thermo Fisher) to which 10% b-mercaptoethanol (Sigma) was added. Samples were then loaded onto a NuPAGE 4-12% Bis-Tris gradient gel (Invitrogen, Thermo Fisher) and blotted onto a nitrocellulose membrane using the iBlot2 dry transfer system (Invitrogen, Thermo Fisher). Membranes were blocked using 5% SureBlock (LubioScience) diluted in 1x PBS containing 0.1% Tween-20 (Sigma Aldrich) for 60 min. Membranes were then incubated with primary antibodies diluted in 1% SureBlock-PBST overnight at 4°C under shaking conditions. For detection, anti-mouse HRP or anti-rabbit HRP (Bio-Rad) were diluted 1:5,000 in 1% SureBlock-PBST. Imaging was performed with a VILBER Fusion Solo S after addition of Immobilon Crescendo Western HRP Substrate (Milipore).

### Primary antibodies

| **Target** | **Manufacturer** | **Dilution** |
| --- | --- | --- |
| GCase | G4171 (Sigma-Aldrich),<br>produced in rabbit | 1: 4,000 |
| Vinculin | ab91459 (Abcam) produced<br>in rabbit | 1:5,000 |
| Beta-Actin (HRP-coupled) | PA1-183-HRP produced in<br>mouse; Invitrogen | 1: 10,000 |
| USF2 | clone 5E9 H00007392-M01<br>(Abnova) produced in rabbit | 1: 1,000 |
| **MITF** | Clone D5G7V #12590<br>(CellSignaling), produced in<br>rabbit | 75.9 ug/mL at 1:1,000 |
| ONECUT2 | 21916-1-AP (Proteintech),<br>produced in rabbit | 0.6 mg/mL at a 1:750 dilution |

### Proteasome inhibitor treatment

For immunoblotting, 250,000 cells per well were seeded in 6-well plates 24 hours before adding MG- 132 (C2211, Sigma-Aldrich) at a concentration of 0.375 µM dissolved in DMSO. 0.375 µM MG-132 containing medium was replaced after 24 hours. Control cells were treated with an equal volume of DMSO. Cells were lysed 48 hours after starting the treatment and immunoblotting for GCase was performed as outlined above.

To assess enzymatic activity, 100,000 cells grown in 96-well plates were treated for 24 hours with either DMSO or 0.5 µM MG-132 in full medium.

On the day of the assay, cells were washed twice in Phenol-red free medium before Hoechst 20 mM (dilution of 1:7,500) was added to three wells per condition, whereas the remaining cells were kept in full medium. Cells were incubated for 20 min at 37°C before imaging of the nuclei with a Pico ImageXpress (Molecular Devices) was performed to assess cell number, as the treatment might result in toxicity/cell death. Thereafter, the enzyme activity assay was assessed in cell lysates as described above. Fluorescence intensities were normalized by the number of nuclei.

The generation of the arrayed CRISPR activation library is outlined in (18)

*The oligonucleotide sequence targeting the TSS of GBA was: sgRNA1 (sgRNA1 sequence) 5’-*

cATGTATGGGTGACAACTTT *-3’; sgRNA2 (sgRNA2 sequence); 5’-* TTGCCCTATAGAGGTGTGTG -3’;

*sgRNA3 5’-* TATAATCTGTAACAGATGAG *-3’; sgRNA4 5’-* GGCACAAGAGGGTGGGACAC *-3’*.

*The oligonucleotide sequence of the NT ctrl was: sgRNA1 (sgRNA1 sequence) 5’-*

GTGTCGTGATGCGTAGACGG *-3’; sgRNA2 (sgRNA2 sequence); 5’-* GTCATCAAGGAGCATTCCGT -3’;

*sgRNA3 5’-* GGACCCTGACATGTATGTAG *-3’; sgRNA4 5’-* GCACTTAGCAGTTTGCAATG *-3’*.

### Lysate-based enzyme activity assay in 96 well plates

The protocol was adapted from a previously published protocol ((56)). Medium was removed by tilting the plates. Cells were washed with 200 µL PBS once before 50 µL of ice-cold McIlvaine buffer (0.1 M citrate and 0.2 M Na2PO4 buffer at pH 5.2 containing 0.4 % Triton-X100 and Protease inhibitor) were added per well. Plates were shaken for 1 h at 4°C at 350 rpm. 50 µL of 4MU-Glc (at a final concentration of 5 mM) in McIlvaine buffer containing 5 mM sodium taurocholate and 1% DMSO was added. Plates were incubated for 1 h at 37 °C before 100 µL of a stop solution (0.3 M Glycine, 0.2 M NaOH solution; pH 10.8) was added.

The plate was centrifuged at 1,000 g for 10 min before fluorescence intensity was read at a VersaMax Microplate Reader (Molecular Devices) or a GloMax® Discover (Promega) (Excitation: 365 nm, Emission: 440 nm).

### Dose-response curves to assess the specificity of the lysate-based assay

4,000 LN-229^WT^ cells were seeded in 96-well plates (ViewPlate-96 Black #6005182, PerkinElmer). The next day, cells were washed in PBS and lysed in 50 mL McIlvaine buffer containing EDTA-free cOmplete Mini Protease Inhibitors (Roche). Cells were shaken at 4°C and 450 rpm for 10 min followed by one hour of incubation at 4°C. Serial dilutions of inhibitors (CBE [575, 383, 256, 170, 114, 76, 50, 33.7 mM final conc.], AT3375 [400, 200, 100, 50, 25, 12.5, 6.25, 3.125 mM final conc.], and miglustat [200, 100, 50, 25, 12.5, 6.25, 3.125, 1.56 mM final conc.]) or an equal volume of DMSO (serving as the negative control) were prepared in an empty 96-well plate in McIlvaine buffer. Cell lysates (three wells per condition) and 4MUG in assay buffer were added (final conc. of 4MUG 2.5 mM) with a multichannel pipette and incubated for one hour at 37°C before the addition of stop solution.

Following a final centrifugation step (300 g for 10 min), plates were read at a VersaMax Microplate Reader (Molecular Devices) (Excitation wavelength: 365 nm, Emission wavelength: 440 nm).

AT3375 was kindly provided by Amicus Therapeutics®.

### Live cell assays to determine GCase, NAGAL and Cathepsin B Activities

The protocols were adapted according to an established protocol obtained from the group of David Vocadlo, Simon Fraser University, Burnaby, British Columbia CA. LysoFQ-GBA and NAGAL-BABS were kindly provided by the group of David Vocadlo (27, 35).

LN-229 cells were plated in 12-well plates (20,000 cells/well). Cells were seeded into 12-well plates. 24 h later, lentiviruses were dispensed manually at an MOI of approximately 4. Cells were collected and counted four days post-transduction and one day before imaging. The cell suspension was plated in 96- or 384-well plates for treatment and imaging (#4680 or 4681, Corning). Dispensing of cells (LN- 229: 1,800 cells/well in 36 μL for 384-well plates or 4,000 cells/well in 90 μL for 96-well plates), with three to six replicates per plate and condition. Plates were centrifuged (150 g for 30 sec) and placed in a humidified incubator. AT3375 was prepared and diluted (in DMEM) and added to the cell-containing plate 24 h before imaging (10 μM final conc.).

On the day of imaging, the substrate solution was prepared in culture medium. For LysoFQ-GBA, the substrate was added to the cell plate at a final conc of 5 μM final. After 1 h of incubation at 37°C, cells were washed three times in Phenol-Red free MEM (Gibco) using a multichannel pipette before 40 or 90 μL imaging buffer was added (Phenol-red free MEM containing 10 % FBS with 1:7,500 dilution of Hoechst and 10 μM AT3375). After another 20 min incubation at 37°C, imaging of live cells was performed using an ImageXpress HT.ai high-content imager (Molecular Devices) connected to environmental control (37 °C, 5% CO_2_). Image acquisition was carried out using a 40x water immersion objective. For each well, 25 (for 384-well plates) to 36 (for 96-well plates) sites (regions of interest) were imaged using the DAPI and FITC channels. Before acquisition, the focus was adjusted for both the DAPI and FITC channels and set to adjust on plate and well bottom. Exposure times were set at 50 ms for the DAPI and 500 ms for the FITC channel.

To assess NAGAL activity, 250 cells/well were seeded in 96-well plates (#4680, Corning) in 90 μL and infected on the following day at an MOI of 4 (assuming a doubling time of 24 h). 24 h before imaging (4 days post-transduction), control wells were treated with the NAGAL-specific inhibitor DGJNAc (10 μM final conc.). On the day of imaging, cells were treated with NAGAL-BABS at a final concentration of the substrate of 10 μM for 3 h. DGJNAc was added to the imaging medium at a final concentration of 10 μM as a negative control.

To assess Cathepsin B activity, the Cathepsin B Assay Kit (Magic Red; Abcam) was used following the protocol of the manufacturer.

### Image analysis

Following acquisition, images were analyzed using the MetaXpress software suite (Molecular Devices). Data were analyzed using the Multiwavelength Cell Scoring module. Nuclei were detected using the DAPI channel and analyzed using a 5/50-μm constraint (min/max width) and a 500 gray-level minimal intensity above background. BODIPY-FL fluorescence was analyzed from the FITC channel image using a 1/7-μm constraint (min/max width) and various minimal intensities above background (200–1,000 gray-level threshold), with the background threshold being set so that there were a few positive pixels in the negative control wells (AT3375- or DGJNAc-treated cells). For each region of interest in each well, this analysis module returned the mean integrated intensity for the FITC or TRITC channel and the number of cells. After combining sites for each well, the integrated intensity normalized by the number of cells for each well was generated and transformed to percentage of activity of the NT ctrl treated cells.

### Bulk RNA Sequencing in LN229^L444P^

High Pure RNA Isolation Kit (Roche) was used for RNA extraction according to the manufacturer’s protocol. Libraries were prepped with the Illumina TruSeq stranded mRNA protocol (Illumina, San Diego, CA, USA) and quality control (QC) was assessed on the Agilent 4200 TapeStation System (Agilent Technologies, Santa Clara, CA, USA). Subsequently, libraries were pooled equimolecular and sequenced on the Illumina NovaSeq6000 platform with single-end 100 bp reads. Sequencing depth was around 20 million reads per sample. Experiments were run in biological triplicates. Data analysis was performed as previously described (57)

### Human induced pluripotent stem cell (iPSC) lines

Human iPSC lines PGPC17, derived from healthy individuals with whole-genome sequencing-based annotation, are generous gifts from Dr. James Ellis at the Hospital for Sick Children, Toronto, Canada (58). Line UOXFi001-B was obtained from EBiSC and was derived from a patient with Parkinson’s Disease (PD) carrying the heterozygous mutation *GBA^N370S/WT^*. The Gaucher Disease (GD) iPSC line C43-1260 (*GBA^L444P/P415R^*) (27) was generated from a GD patient’s skin fibroblast line (GM01260, Coriell) and characterized using a contract service provided by the Tissue and Disease Modelling Core (TDMC) at the BC Children’s Hospital Research Institute following their standard protocols. The genomic integrity of this line was confirmed by G-banded karyotyping performed at WiCell. The pluripotency of C43-1260 was tested by in vitro differentiation to the three germ layers using the STEMdiff™ Trilineage Differentiation Kit (STEMCELL Tech, Catalog # 05230). All iPSCs were cultured and expanded in mTeSR™ Plus medium (STEMCELL Tech, Catalog # 100-0276) according to the manual.

### Generation of iPSC-derived neural progenitor cells (NPCs) and forebrain mature neurons

iPSC lines including PGPC17, UOXFi001-B, and C43-1260 were differentiated to NPCs using the STEMdiff™ Neural Induction Medium + SMADi (STEMCELL Tech, Catalog # 08581) following the monolayer-based differentiation protocol described in the technical manual. Generated NPCs were either used directly or cryopreserved in STEMdiff™ Neural Progenitor Freezing Medium (STEMCELL Tech, Catalog # 05838) and stored in a liquid nitrogen Dewar until use.

NPCs were further differentiated to neuronal precursors using the STEMdiffTM Forebrain Neuron Differentiation Kit (STEMCELL Tech, Catalog # 08600) and then matured to forebrain neurons using the STEMdiffTM Forebrain Maturation Kit (STEMCELL Tech, Catalog # 08605) following the standard protocol outlined in the technical manual.

### Live-cell GCase assay on iPSC-derived forebrain mature neurons transduced with lentiviral vectors

Neuronal precursors were seeded in 96-well High Content Imaging (HCI) plates (Corning, Catalog # 4680) precoated with poly-L-ornithine (PLO, 15 μg/mL) and laminin (5 μg/mL) at a seeding density of approximately 100,000 cells/cm2 and cultured using the Forebrain Maturation Kit as described above. Half-medium changes were performed every 2 - 3 days. Seven days post-seeding, neurons were infected with an appropriate amount of lentiviruses encoding different transcription factors or with lentiviral vectors encoding GBA isoform two or an empty vector serving as positive and negative controls, respectively. The following day, an 80% medium exchange was carried out to reduce toxicity related to lentiviral particles and to minimize the detachment of neurons, followed by an additional 80% medium exchange 48 hours later before returning to half medium exchanges every 2 - 3 days.

Live-cell assays to assess GCase activity using the fluorescence-quenched probe LysoFQ-GBA (27) were carried out on days 7 and 14 post-infection. The assay was initiated by replacing the culture medium with fresh medium containing 10 µM of LysoFQ-GBA and then incubating in the TC incubator for designated lengths of time (2 h for the GBA WT line PGPC1, 3 h for the PD-GBA line UOXFi001-B, and 4 h for the GD line C43-1260). The assay was stopped by two consecutive 85% medium exchanges with BrainPhys™ Imaging Optimized Medium (IOM) (STEMCELL Tech, Catalog # 05796) and one 85% medium exchange with IOM supplemented with NeuroCult™ SM1 Neuronal Supplement (SM1) (STEMCELL Tech, Catalog # 05711) containing 10 μM of AT3375. After 20 min of incubation at 37°C, 10 μL of 11 x Hoechst (11 μg/mL) prepared in IOM + SM1 were added to each well (final conc 1 μg/mL). The plate was gently tapped for distribution, centrifuged at 300 x g for 30 s, and incubated for another 10 min. Images were then acquired using a high-content widefield microscope (ImageXpress XLS, Molecular Devices) with a 40x objective. Micrographs were analyzed using the in-built HCI analysis software MetaXpress (Molecular Devices) to yield the mean integrated intensity level of vesicle-like structures (representing lysosomal GCase activity) per cell in the FITC channel for each well.

To assess neuron quality using immunofluorescence, forebrain neurons cultured in a 96-well HCI plate were fixed with 4% PFA in PBS for 20 min at RT and permeabilized using 0.1% Triton X-100 in PBS for 5 min on ice. After blocking with 5% BSA in PBS, neurons were incubated with primary anti-GFAP (Aves Lab, SKU: GFAP) or anti-MAP2 (Abcam, Catalog # Ab5392) antibodies at 4°C overnight. On the following day, primary antibodies were washed away, and fixed neurons were stained with corresponding secondary fluorescent antibodies. Neurons were finally stained with Hoechst for 30 min and imaged using the ImageXpress XLS high-content widefield microscope (Molecular Devices).

### Proteomics analysis

LN-229^L444P^ dCas9-VPR cells transduced with qgRNAs targeting *ONECUT2* or the non-targeting control at an MOI of 4 were prepared in four biological replicates (separate wells of a 6-well plate), washed twice in PBS, harvested by scraping of cells in PBS, and snap-frozen in liquid nitrogen. Native proteins were extracted by resuspending the pellets in a buffer (100 mM HEPES, 150 mM KCl, 1 mM MgCl2, pH 7.4), and lysed with 10 cycles of 10 dounces each using a pellet pestle on ice. Protein concentrations were measured with bicinchoninic acid assay and adjusted. Samples were boiled for 5 minutes at 99°C and cooled down on ice. The samples were diluted in a 10 % sodium deoxycholate to a final DOC concentration of 5 %, reduced with tris(2-carboxyethyl)phosphine hydrochloride (TCEP) in a final concentration of 5 mM at 37 °C for 40 minutes, and alkylated using a final concentration of 40 mM iodoacetamide (IAA) for 30 minutes at room temperature in the dark. The samples were then diluted to 1 % DOC using 10 mM ammonium bicarbonate. Proteins were digested overnight at 37 °C with LysC (Wako Chemicals) and trypsin (Promega) in a 1/100 enzyme-to-substrate ratio, under constant shaking. Formic acid (FA) (Carl Roth GmbH) was added at a final concentration of 2 % to stop digestion and precipitate DOC. DOC was removed by three centrifugation cycles, and subsequently, the supernatant was desalted on Sep-Pak tC18 cartridges (Waters). Samples were eluted with 50% acetonitrile and 0.1% formic acid. LC-MS/MS experiments were performed on an Orbitrap Eclipse mass spectrometer (Thermo Fisher Scientific). Data-independent acquisition (DIA, ref) scans were performed in 41 variable-width isolation windows. MS data were searched with Spectronaut (Biognosis AG). Statistical analysis was performed in R.

Ref: Mol Cell Proteomics 11, O111 016717. 10.1074/mcp.O111.016717.

---

**Supplementary Fig. 1.**
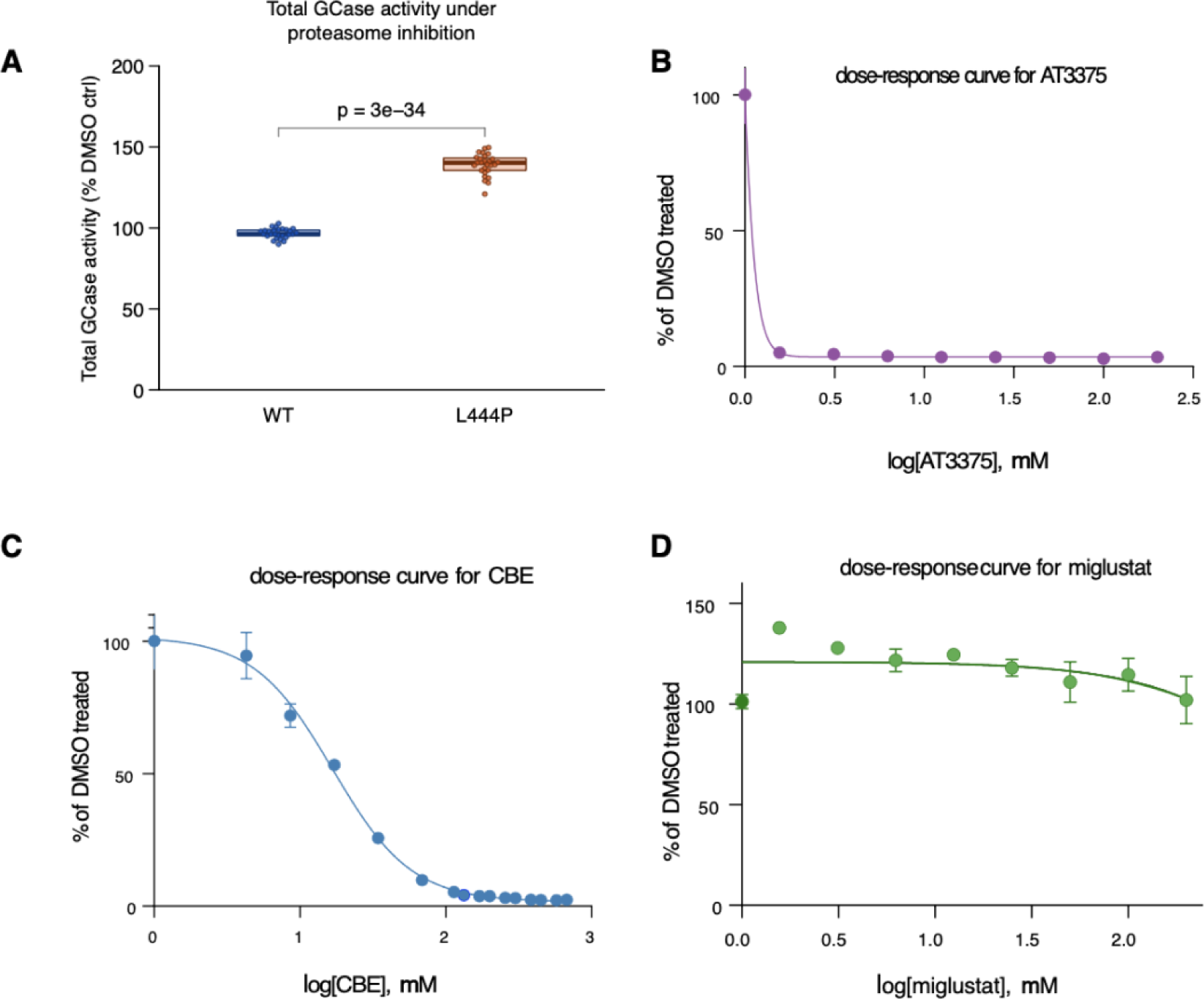
Validation of the lysate-based assay employing 4MU-Glc to quantify total GCase activity in LN-229^WT^ cells. A. Lysate-based assay in LN-229^WT^ and LN-229^L444P^ after treatment with MG-132 (N=2 plates). B. Dose- response curve for conduritol-beta-epoxide (CBE). C. Dose-response curve for AT3375, a highly selective GCase inhibitor, in LN-229^WT^ cells. D. Dose-response curve for miglustat, a GBA2 inhibitor (N=1 experiment per inhibitor).

**Supplementary Figure 2.**
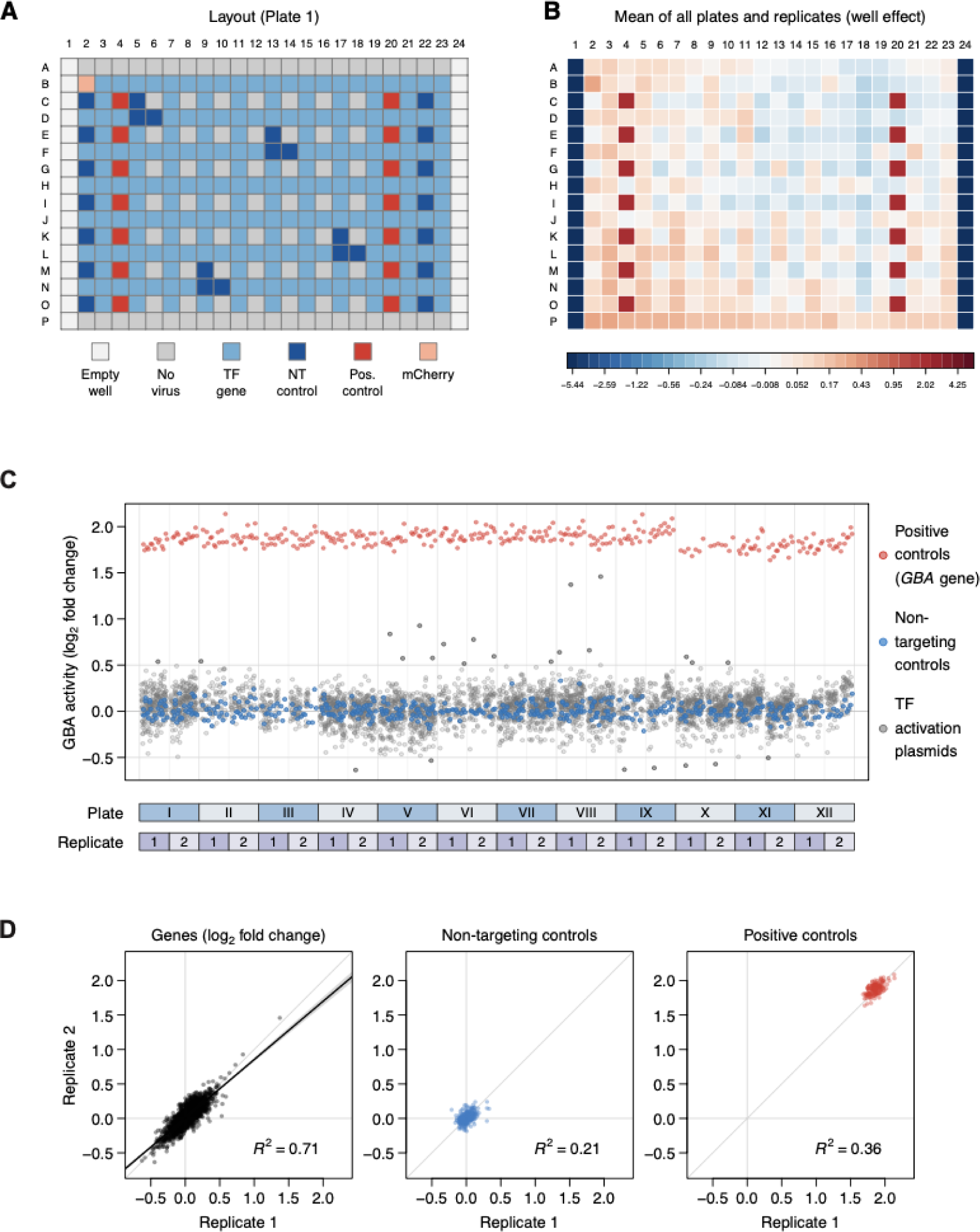
Quality metrics of primary arrayed CRISPRa screen. A. Pipetting scheme showing plate layout adopted in the primary screen with distribution of controls and TF genes. Outermost wells were left untransduced. B. Heat map depicting mean fluorescence intensity of all plates and replicates. C. Plate-well series plot of all 24 assay plates on which enzymatic activity was assessed. D. Replicate correlation of all samples and control populations with Pearson correlation coefficient (R^2^).

**Supplementary Fig. 3.**
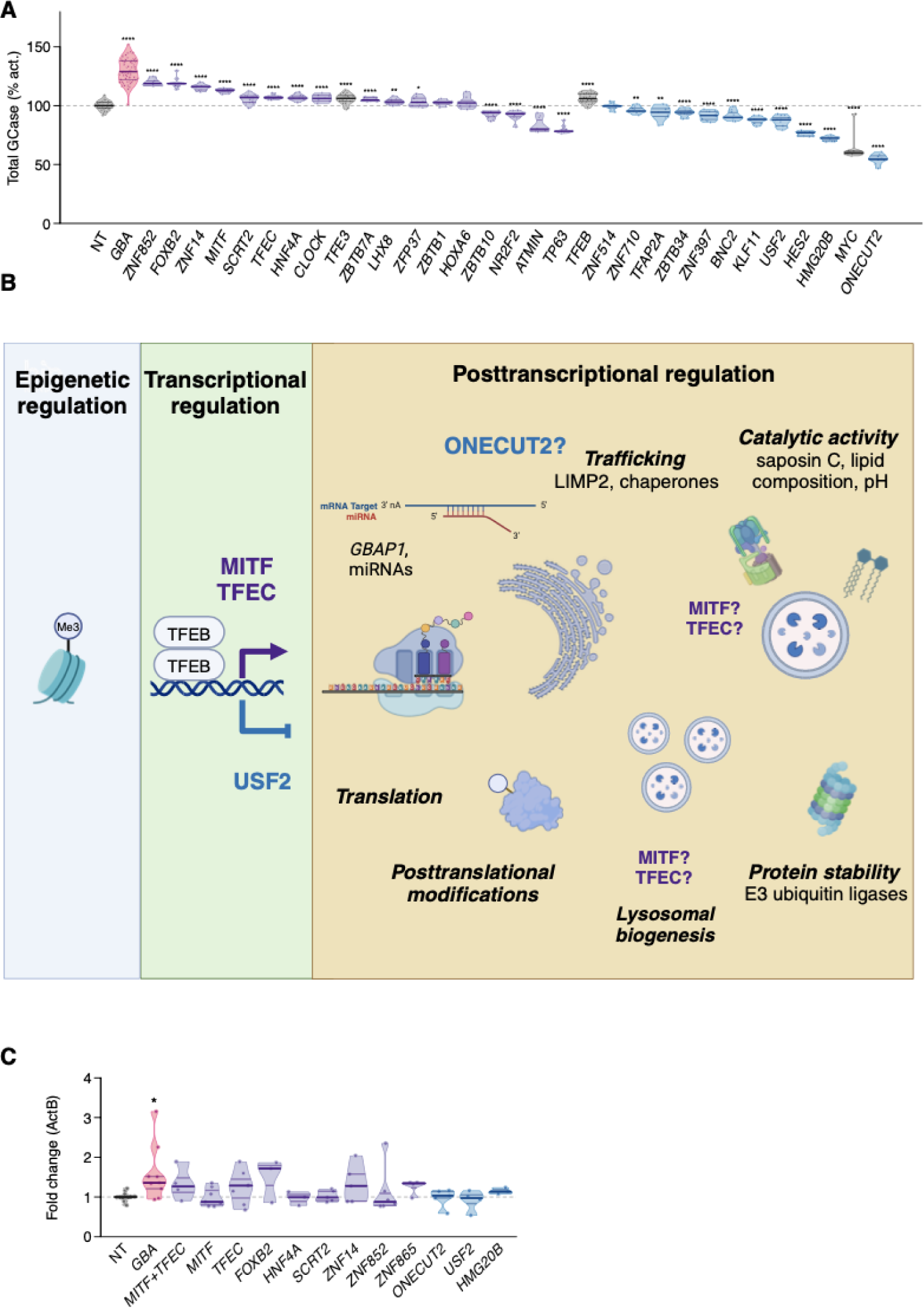
A. Testing of identified candidates in LN-229^WT^ in cell lysates. NT ctrls and TFs previously described to regulate GCase expression (TFE3, TFEB, and MYC) are shown in grey. TFs that increased fluorescence intensity in the primary screen as compared to NT ctrls in LN-229^L444P^ are shown in purple, TFs decreasing enzymatic activity are colored in blue, positive ctrls (GBA-activated cells) are depicted in red. Abscissa: Total GCase activity measured in cell lysates and expressed as % activation, with median fluorescence values of the NT ctrl wells set as 100% (N >= 6 transductions in >=1 experiment). A Welch’s t-test was performed to assess ps. **** p<0.0001; *** p<0.001; ** p<0.01; * p<0.05. B. Layers of regulation of GBA expression and GCase activity and hypothesized regulatory mechanisms of selected candidate TFs. C. Fold changes in RNA levels of GBAP1 assessed by RT-qPCR following CRISPRa of the candidate TFs designated on the x-axis. Actin beta was used as the reference (N = 3-8 repeats).

**Supplementary Figure 4.**
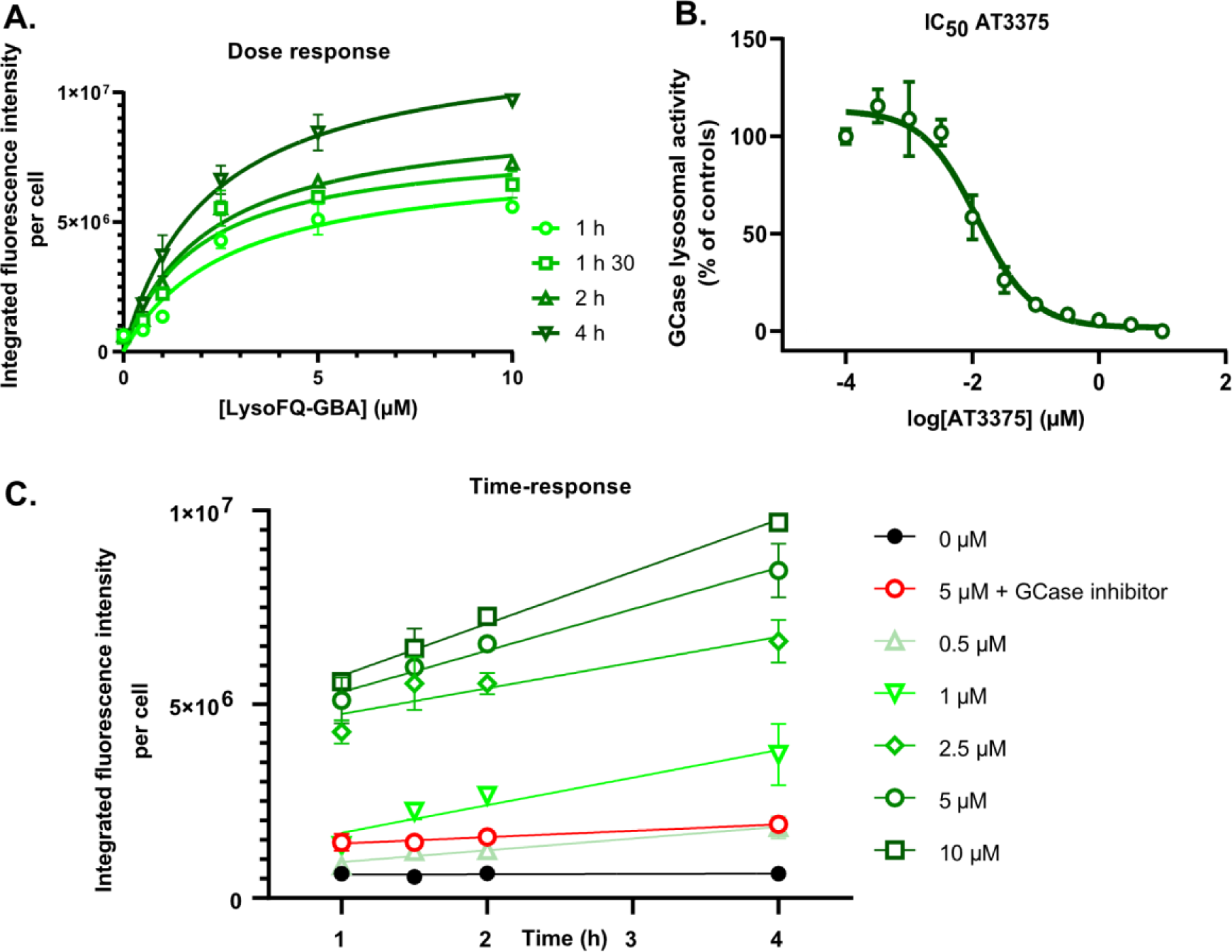
Dose (A)- and time (C)-response curves for LysoFQ-GBA in LN-229^WT^ cells. Integrated fluorescence intensity was averaged from region of interest per well and normalized to the number of nuclei. 10 μM AT3375 was used as a specific GCase inhibitor at 10 μM in panel C to confirm the specificity of the fluorescence signal for GCase. B. IC50 curve for AT3375. Cells were treated with a range concentration of AT3375 (100 pM -10 μM) for 5 hours and subsequently incubated with 2.5 μM of LysoFQ-GBA for 1 hour. Vehicle (DMSO)-treated cells served as a reference for GCase activity normalization. A-C: N= 3 independent wells per condition

**Supplementary Figure 5.**
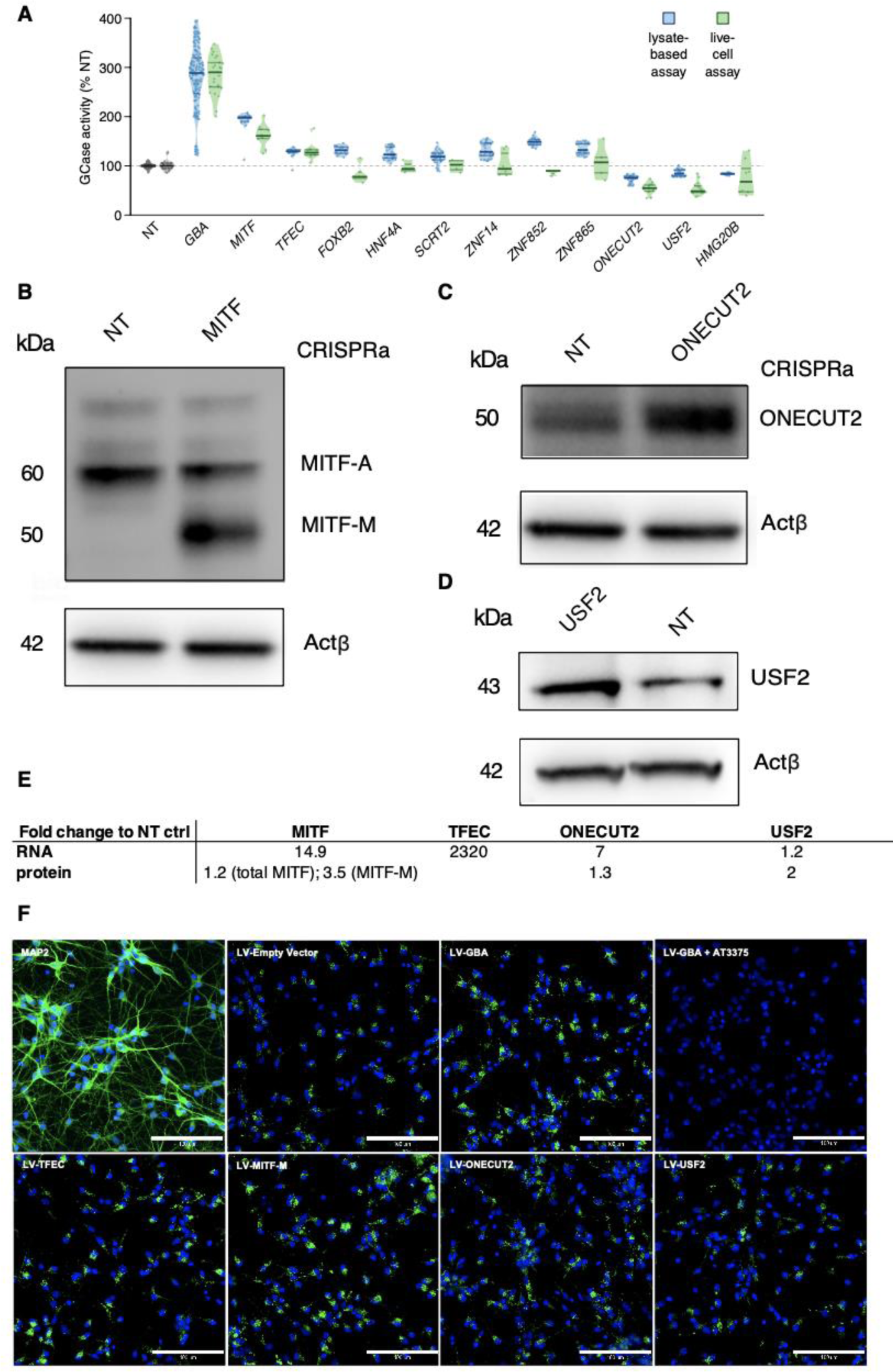
A. Comparison of results of CRISPRa of candidates on GCase activity between the lysate-based (4MUG-Glc; blue) and the live-cell assay (LysoFQ-GBA; green) in LN-229^L444P^. B. CRISPRa of the proximal transcription start site of *MITF* predominantly increases the shortest MITF isoform (MITF-M) lacking the mTORC1 phosphorylation site. Total MITF increases by 120% as compared to the NT ctrl infected LN-229^L444P^ dCas9-VPR cells, whereas MITF-M increases by 350% (N = 1 experiment). C. CRISPRa of *ONECUT2* results in a 130% increase in the protein expression levels as compared to the NT ctrl infected LN-229^L444P^ dCas9-VPR cells. D. CRISPRa of *USF2* results in a 200% increase in protein expression levels as compared to the NT ctrl infected LN-229^L444P^ dCas9-VPR cells. E Fold changes following CRISPRa of candidate TFs after lentiviral transduction (MOI = 3- 4) in LN-229^L444P^ dCas9-VPR cells at RNA and protein level as compared to NT infected cells (N =1 experiment). F. Representative images of MAP2, a neuron-specific cytoskeleton marker, and the turnover of LysoFQ-GBA after treatment with lentiviral (LV) overexpression vectors encoding indicated candidate TFs 14 days post-transduction in iPSC-derived forebrain neurons from a PD patient with a heterozygous *GBA^N370S/WT^* mutation. AT3375 = GCase specific inhibitor (Scale bars: 100 μm).

**Supplementary Fig. 6.**
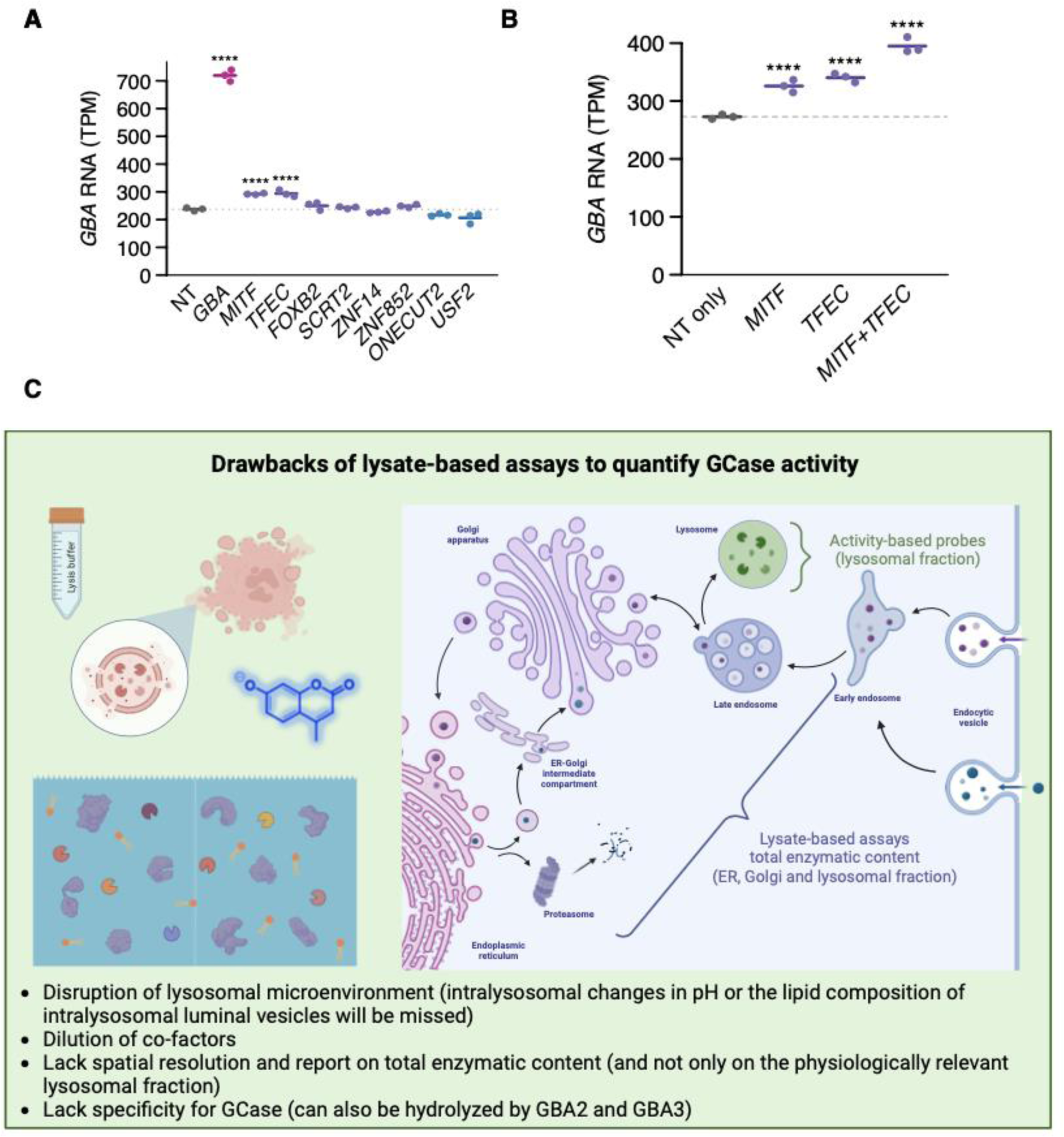
A-B. Bulk RNA Sequencing data depicting transcripts per million (TPM) of *GBA* RNA levels after CRISPRa of designated candidate TFs (N = 3 replicates). C. Schematic summarizing major drawbacks of lysate-based assays. **** p<0.0001;*** p<0.001; ** p<0.01; * p<0.05

**Supplementary Fig. 7.**
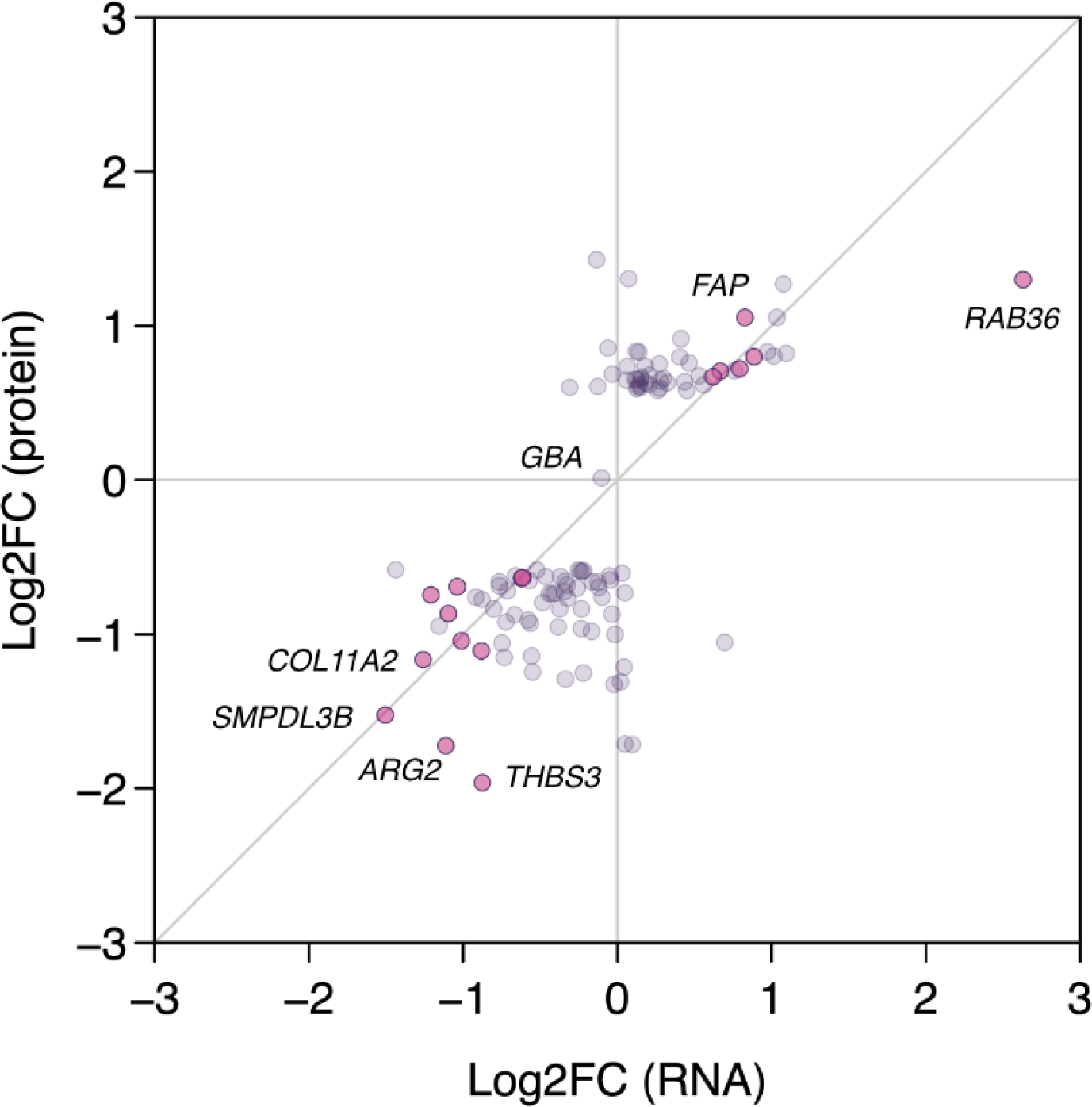
Comparison of differentially expressed genes identified in bulk RNASeq (x-axis) with protein abundances (y-axis) identified by mass spectrometry following CRISPRa of *ONECUT2* in LN-229^L444P^ dCas9-VPR cells. Differentially expressed genes reaching a Log2FC ≥ 1.5 and a false-discovery rate adjusted p < 0.05 at both the RNA and protein level are highlighted in pink. N = 3 replicates for bulk RNASeq and N = 4 replicates for mass spectrometry analysis

**Supplementary Fig. 8.**
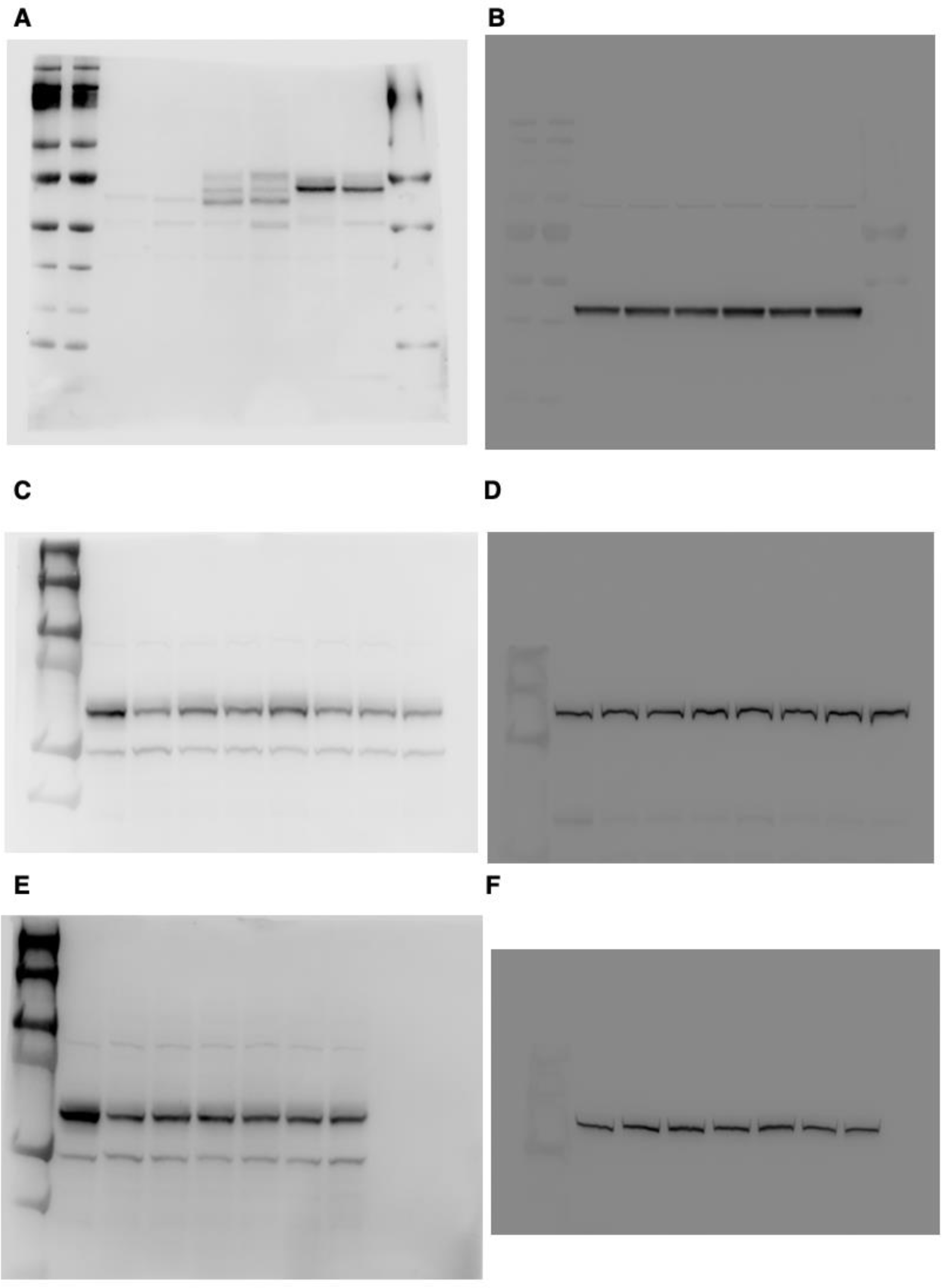
Original, uncropped Western blots of Fig. 1D showing GCase (A.) and beta-Actin (B.). Fig. 3C for GCase (C and E) and Vinculin (D and F) in LN-229^L444P^ dCas9-VPR cells.

**Supplementary Fig. 9.**
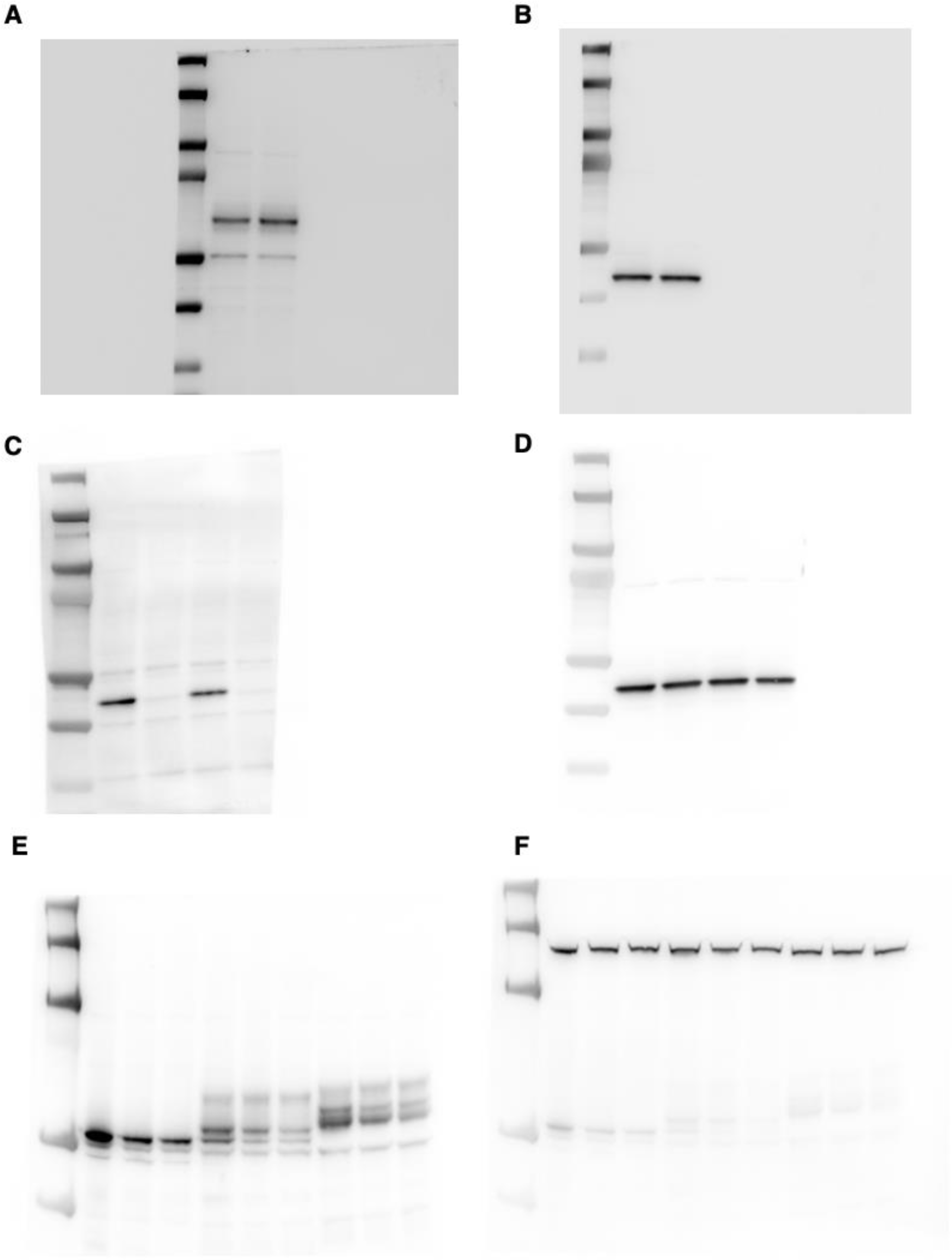
Original, uncropped Western blots of Fig. 4C showing GCase and beta-Actin levels (A-B.) in LN-229^L444P^ Cas9 cells. Fig. 4E showing USF2 and beta-actin levels in LN-229^L444P^ Cas9 cells (C-D.) and Fig. 4G showing GCase and beta-Actin levels in LN-229^WT^ Cas9 cells (E-F.).

